# BioMINTREL expands biological problem solving through cross-disciplinary representations

**DOI:** 10.64898/2026.10.10.757703

**Authors:** Jiaqi Dong, Yilan Zhang, Elva Gao, Changchun Yang, Jiacheng Wang, Chao Zha, Zhenyu Wu, Xiaopeng Xu, Juexiao Zhou, Xin Gao

## Abstract

Biological research agents often rely on familiar analytical methods, limiting the questions explored in existing data. We introduce BioMINTREL, an AI agent that finds methods from other disciplines and connects them to biological measurements. It can draw on relationships among datasets and complementary ideas from several methods to develop new questions and analyses. We evaluate it on BioMDC-Bench, which assesses 228 possible applications of computational methods to biological tasks. Compared with GPT-5.4 mini and Qwen3.6-35B, BioMINTREL improves F1 agreement with human experts by up to 45.3 percentage points and increases coverage of applications supported by studies published in subsequent years by up to 58.8 percentage points. Four case studies on real data go beyond previous analyses to reveal dose-specific variant effects, constraints on spatial cell communication and routes through perturbation-response states, and identify the sampling needed to test whether rare high-state cells predict later lineage size. These results highlight BioMINTREL’s potential to expand questions in existing data and suggest directions for subsequent research.

## Introduction

The growing scale and diversity of biological data create new opportunities for discovery and increase the need for computational approaches that translate complex measurements into biological understanding[1–3]. Recent advances in large language models (LLMs) have enabled agents to participate in literature search, hypothesis generation, data analysis and interpretation[4–6]. However, broader automation does not necessarily expand the range of scientific questions that researchers consider. In biology, the questions asked of existing data may still be shaped by familiar concepts and analytical methods within the field. Researchers and autoresearch systems that rely on familiar analytical frameworks may therefore overlook questions that different ways of organizing the same measurements could make accessible.

The history of computational biology shows that a new mathematical representation can itself change the questions that data can answer. A representation organizes measurements into objects and relationships that a method can analyse. For example, traditional trajectory methods can order cells by similarity in transcriptional state. Waddington-OT used optimal transport, a method for matching distributions, to connect cell populations at different times through probabilistic links. These links enabled inference of ancestor–descendant relationships and revealed developmental paths and regulators that are difficult to recover from static similarity alone[7]. Hidden Markov models organize genomic sequences as probabilistic state processes across different functional regions and enable joint prediction of complete gene structures[8]. de Bruijn graphs reframe short-read assembly as a path problem on a graph[9, 10]. In these examples, the biological measurements did not change. The mathematical representation changed, along with the relationships that could be inferred. Established methods from other fields can therefore make more biological questions accessible in existing data.

Recent AI systems have expanded support for biological research across analysis planning, tool use and interpretation. Robin and the Virtual Lab coordinate scientific reasoning and validation[11, 12]; CellVoyager proposes and implements single-cell analyses[13]; Biomni connects biomedical tasks to tools and databases[14]; and BiOmics supports the integrated interpretation of multiple types of molecular data[15]. These systems mainly use existing biomedical tools, models and analytical frameworks, with limited systematic exploration of methods and data representations from outside biology. Cross-domain analogical reasoning uses structural similarities across scientific fields to generate more diverse and novel candidate solutions for a predefined research problem[16]. This raises an earlier question: when the research question has not yet been fixed, can methods from other fields help determine what else an existing biological dataset can reveal? Answering this question requires defining the method’s objects, relationships and computational operations from the available measurements, so that the data can guide which research directions to develop.

Here we introduce BioMINTREL (**Bio**medical **M**ethod **In**ference and **T**ransfer from **R**esearch **E**vidence and **L**iterature), an AI agent that expands the problem-solving space available to biomedical agents: the questions and methods they can consider for available data. BioMINTREL draws on established methods from other disciplines and can start from biological data without a predefined question. It can formulate a joint task from relationships among several datasets, or develop a model by relating the computations and ideas of several methods. It also draws on information from scientific text, figures, tables and supplementary materials to define analysis inputs and interpret results. Scientific fields often use different terms to describe similar analytical structures[17, 18], so BioMINTREL searches using both the scientific setting and the mathematical structure of a problem. It then assesses method–data correspondence, which defines how a method’s required objects and relationships can be constructed from the available measurements. This correspondence also specifies how the core computation addresses a biological question. These definitions guide biological questions and analysis plans that the data can support. BioMINTREL then executes selected plans with comparisons and controls to test the representation, interpret the results and identify directions for further study.

Across GPT-5.4 mini[19] and Qwen3.6-35B[20], BioMINTREL agrees more closely with experts than direct judgments from the same models. A retrospective analysis also shows that it identifies more applications supported by later literature. We then apply BioMINTREL to four real-data case studies covering protein variants, spatial cell communication, perturbation responses and cell lineages. These analyses distinguish effects across drug doses, separate local constraints on communication and compare perturbation-response routes. They also identify the sampling needed to test whether a few cells with high expression of selected gene sets predict later lineage size beyond cell number and average state. Our search of studies published through 31 August 2025, the knowledge cutoff of GPT-5.4 mini, found no prior work matching these four specific analyses. Together, these results show how cross-disciplinary methods can broaden questions in existing data and guide further investigation.

## Results

BioMINTREL starts from one or more biological data papers and one or more method papers from other disciplines (Fig. 1a). A task combines the selected data with a question supplied by the user or identified from the data. With several datasets, BioMINTREL examines complementary measurements or links between biological stages to define a joint task. With several methods, it considers complementary computations, successive steps and suitable comparisons when formulating the analysis. Candidate methods are ranked using scientific context, which describes their research settings and applications, and analytical structure, which describes their mathematical objects, relationships and computational requirements. BioMINTREL assesses method–data correspondence for each task–method pair and prioritizes supported applications. It then develops complete analysis plans specifying the biological question, required calculations, comparisons and interpretation. Selected plans are compared with prior work, and those taken forward are executed with the planned controls. The results and related literature guide biological interpretations and follow-up questions (Methods).

**Figure 1:**
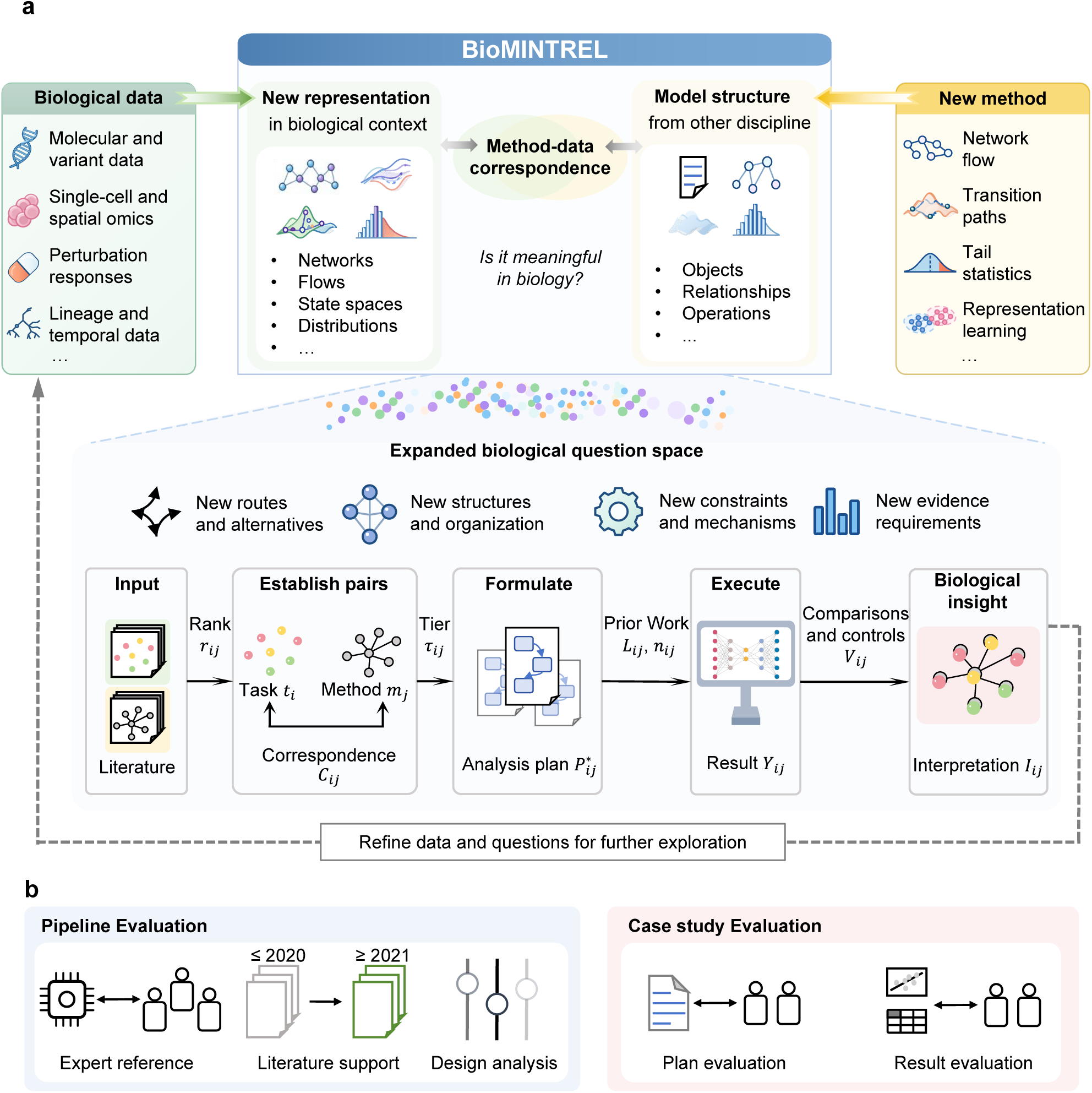
BioMINTREL expands biological question space. **a**, Methods from other disciplines provide new representations of biological data. BioMINTREL defines biological counterparts for the objects, relationships and operations required by each method. Starting from data and method papers, it ranks candidate methods using scientific context and analytical structure, summarized by the retrieval score *r_ij_*. Each task *t_i_* combines a dataset with a starting biological question, and *m_j_* denotes a candidate method. The method–data correspondence *C_ij_* defines how the method applies to the data and which question it can address. The adaptation tier *τ_ij_* guides selection of candidate pairs for complete analysis plans 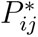. Before execution, BioMINTREL searches related studies *L_ij_* and assesses the plan’s relation to prior work *n_ij_*. Analysis results *Y_ij_*, comparisons and uncertainty estimates *V_ij_*, and relevant literature support the biological interpretation *I_ij_*, which guides further measurements and questions. **b**, Pipeline evaluation compares BioMINTREL’s judgments with expert consensus based on literature through 2020 and with related studies published from 2021 onward. In four case studies, experts first compare analysis plans without knowing their sources and then assess the completed analyses and supporting evidence.

We evaluated BioMINTREL at three levels (Fig. 1b). First, we constructed BioMDC-Bench (Benchmark for Method–Data Correspondence in Biology), comprising 228 pairs of 19 biological tasks and 12 candidate methods. Each pair has a unanimous judgment from three experts based on literature through 2020 and a separate record of whether related method applications were reported from 2021 onward (Evaluation 1.1; Evaluation 1.2). Second, we varied how candidate methods were found and ranked, and tested whether generating several independent plans increased the chance of obtaining a complete plan. Third, experts evaluated both the proposed plans and completed analyses for four case studies on real biological data.

### 1 Expert consensus and later literature jointly support BioMINTREL in identifying more method applications

#### 1.1 BioMINTREL is closer to expert consensus than direct LLM judgment

Across the 228 task–method pairs in BioMDC-Bench, BioMINTREL’s judgments are closer to the expert reference than direct LLM judgments. T1 denotes direct application, T2 denotes application with defined adaptations that retain the method’s core computation, and T3 denotes a correspondence that has not been established. We evaluate T1+T2 against T3. The direct LLM baseline uses the same underlying model and task and method descriptions to assign these categories without explicitly constructing the method–data correspondence (Method 3.2; Evaluation 1.1). We use F1 to summarize how accurately and completely each approach identifies the expert-supported T1+T2 pairs. BioMINTREL increases F1 from 65.2% to 84.7% with GPT-5.4 mini and from 34.0% to 79.3% with Qwen3.6-35B, gains of 19.5 and 45.3 percentage points, respectively.

BioMINTREL identifies more pairs for which experts consider the method–data correspondence established. Among the 176 pairs assigned to T1+T2 by experts, BioMINTREL identifies 150/176 (85.2%), compared with 90/176 (51.1%) for GPT-5.4 mini. BioMINTREL identifies 134/176 (76.1%), compared with 36/176 (20.5%) for Qwen3.6-35B. Across all 228 pairs, BioMINTREL agrees with expert judgments more often than direct judgment by the same model (two-sided exact McNemar tests, *P* = 1.38 × 10*^−^*^5^ and 1.42 × 10*^−^*^9^, respectively; Fig. 2a; Evaluation 1.1.2).

**Figure 2:**
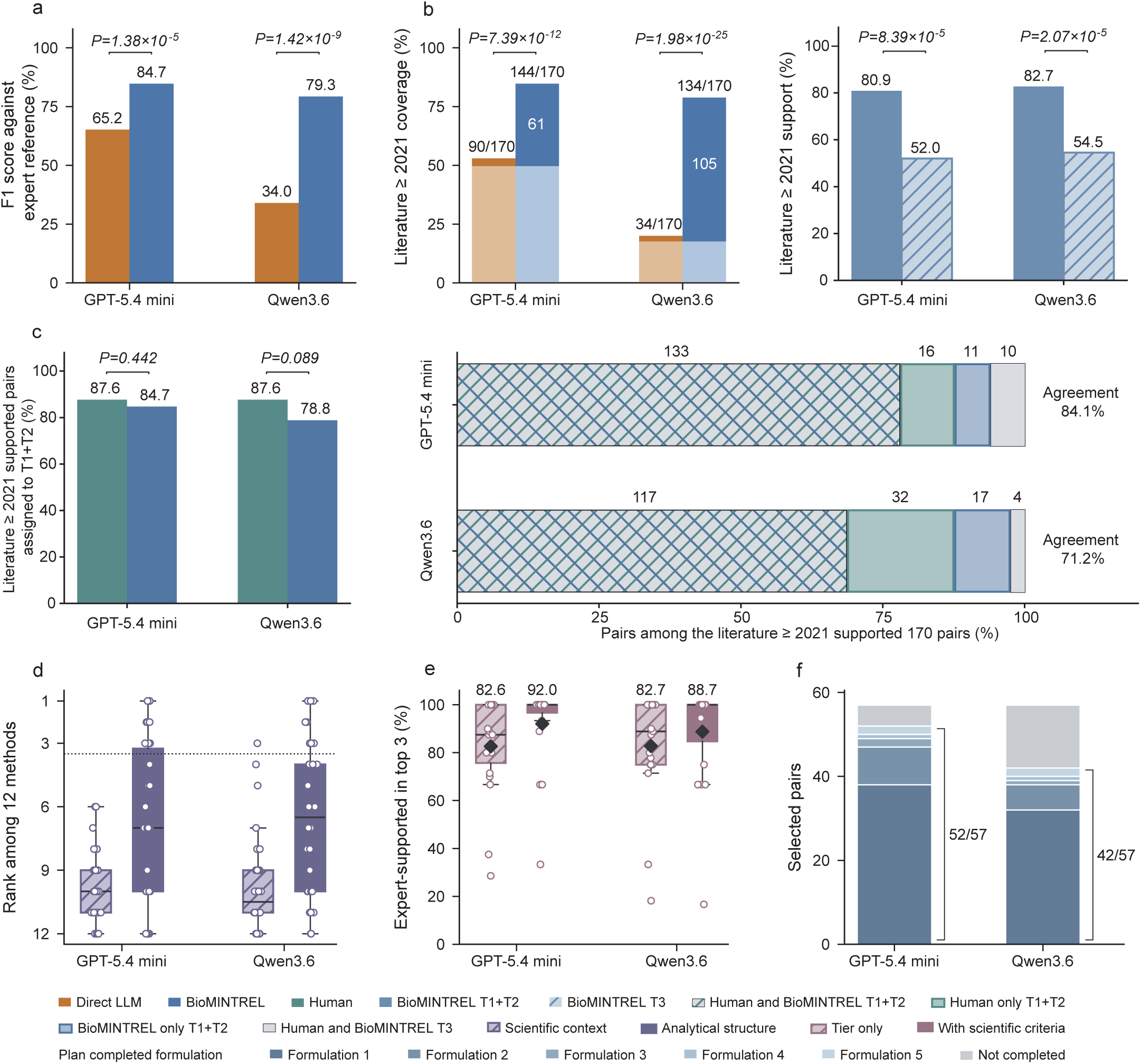
BioMINTREL identifies supported method–data correspondences and develops them into complete analysis plans. **a**, F1 scores against the expert reference across 228 task–method pairs for direct LLM judgment and BioMINTREL. The two-sided exact McNemar tests compare paired correctness against the expert reference. **b**, Left, coverage of the 170 pairs supported by literature published from 2021 onward; darker segments indicate pairs identified only by the corresponding approach. Right, later-literature support among pairs assigned to T1+T2 or T3 by BioMINTREL. **c**, Human and BioMINTREL T1+T2 coverage among the same 170 later-supported pairs, with counts of their joint binary judgments. Agreement counts pairs with the same T1+T2 or T3 assignment. **d**, Ranks of 38 expert-reference T2 methods with the lowest Scientific context similarity under Scientific context and Analytical structure retrieval. The dotted line marks the top-three boundary. **e**, Expert-supported proportion among the three highest-priority methods across 19 tasks using Tier alone or Tier together with the three scientific criteria. Boxes show the median and interquartile range, with whiskers extending to 1.5× the interquartile range. **f**, First complete analysis plan across five independent Formulations for 57 selected task–method pairs; grey indicates pairs with no complete plan among the five candidates.

#### 1.2 BioMINTREL covers more pairs for which related research later appeared

Related studies published from 2021 onward provide a second, retrospective evaluation of method applications. These studies use the same or related methods for the target task or a related biological question (Evaluation 1.2). Among the 170 pairs with later-literature support, BioMINTREL increases T1+T2 coverage from 90/170 (52.9%) to 144/170 (84.7%) with GPT-5.4 mini and from 34/170 (20.0%) to 134/170 (78.8%) with Qwen3.6-35B. The paired comparison identifies 61 pairs selected only by BioMINTREL versus 7 only by direct GPT-5.4 mini, and 105 versus 5 with Qwen3.6-35B (two-sided exact McNemar tests, *P* = 7.39 × 10*^−^*^12^ and 1.98 × 10*^−^*^25^, respectively; Fig. 2b, left).

Across all 228 pairs, BioMINTREL expands the T1+T2 sets from 100 to 178 pairs with GPT-5.4 mini and from 36 to 162 with Qwen3.6-35B. Later-literature support covers 90/100 (90.0%) pairs selected by direct GPT-5.4 mini and 144/178 (80.9%) selected by BioMINTREL. For Qwen3.6-35B, the corresponding numbers are 34/36 (94.4%) and 134/162 (82.7%). BioMINTREL thus expands the range of task–method pairs available for further study and identifies applications missed by the direct LLM but supported by later research (Evaluation 1.2.2).

Pairs classified as T1+T2 by BioMINTREL have a higher rate of later-literature support than those classified as T3. With GPT-5.4 mini, later-literature support is 80.9% for T1+T2 and 52.0% for T3. With Qwen3.6-35B, it is 82.7% for T1+T2 and 54.5% for T3 (two-sided Fisher exact tests, *P* = 8.39 × 10*^−^*^5^ and 2.07 × 10*^−^*^5^, respectively; Fig. 2b, right; Evaluation 1.2.2).

Among the same 170 pairs with later-literature support, experts assign 149/170 (87.6%) to T1+T2, compared with 144/170 (84.7%) for BioMINTREL (GPT-5.4 mini) and 134/170 (78.8%) for BioMINTREL (Qwen3.6-35B). Expert and BioMINTREL judgments agree on 143/170 (84.1%) and 121/170 (71.2%) pairs, respectively. After Holm correction for the two comparisons, paired coverage tests give *P* = 0.442 and 0.089, respectively (two-sided exact McNemar tests; Fig. 2c; Supplementary Note S3.4).

### 2 Retrieval representations, scientific criteria, and repeated generation improve candidate discovery and plan development, respectively

We further compare settings for finding methods, ranking candidates and generating analysis plans in BioMINTREL using either GPT-5.4 mini or Qwen3.6-35B as the underlying language model (see Evaluation 2).

#### 2.1 Analytical structure brings methods ranked low by context retrieval into candidate comparison earlier

For each model, we examine the 38 expert-reference T2 pairs with the lowest scientific context similarity. We compare how their methods rank among the 12 candidates for each task when retrieval uses scientific context or analytical structure alone. With GPT-5.4 mini, analytical structure increases the number in the top three from 0/38 to 10/38, and the median rank moves from 10th to 7th. With Qwen3.6-35B, the number increases from 1/38 to 8/38, and the median rank moves from 10.5 to 6.5. Analytical structure thus brings methods with different original application settings into consideration (Fig. 2d; Evaluation 2.1).

#### 2.2 The three scientific criteria increase the number of expert-supported methods in the top three

Each task selects three methods to continue to analysis-plan generation. Ranking therefore determines which analysis directions are developed first. The three scientific criteria assess the correspondence’s biological basis, the contribution of the method’s core computation to the main result and whether the planned comparisons can independently test that result. With the assigned categories unchanged, adding these criteria increases the mean proportion of expert-supported methods among the top three from 82.6% to 92.0% with GPT-5.4 mini and from 82.7% to 88.7% with Qwen3.6-35B. Uniform selection of three methods gives an expected proportion of 77.2%. These proportions account for ties at third place using the fraction of expert-supported methods among the tied candidates (Fig. 2e; Method 3.3; Evaluation 2.2).

#### 2.3 Five independent generations give more pairs complete plans

Five independent generations give more selected task–method pairs at least one complete analysis plan. A complete plan specifies the data inputs, biological question, core computation, analysis steps and comparisons needed to test the result (Method 4.2). With GPT-5.4 mini, 38/57 pairs obtain a complete plan after the first generation, increasing to 52/57 after five independent generations (66.7% to 91.2%). With Qwen3.6-35B, the number increases from 32/57 to 42/57 (56.1% to 73.7%; Fig. 2f; Evaluation 2.3).

For both models, the number of tasks with at least one complete plan increases from 18/19 after the first generation to 19/19 after five independent generations.

### 3 Expert evaluation supports the plans’ new analytical value and further research

Two experts compare three plans for each of four cases without knowing their sources: the BioMINTREL plan, a plan using established methods in the field and one using conventional statistical comparisons. All three use the same data and review question (Evaluation 3.1). The mean analytical novelty score relative to the original paper is 3.38/4 for BioMINTREL and 2.50/4 for both comparison plans. Across the eight reviews, experts choose BioMINTREL as most likely to provide additional information five times, rank it first as most worth executing four times, including one tied first place, and recommend proceeding with it seven times (Supplementary Data 3 (`plan-review forms`)).

Experts recommend further research in seven of eight assessments of the completed BioMINTREL analyses. Mean scores are 3.63/4 for biological relevance, 3.50/4 for the match between the analysis framework and the question, and 3.13/4 for information gain relative to conventional analysis (Evaluation 3.2). Suggestions include testing the stability of candidate mutations across thresholds, validating local spatial relationships, comparing graph constructions and testing lineage predictions with matched cell counts. BioMINTREL’s follow-up proposals identify the same priorities with human experts for spatial validation, graph comparisons and lineage tests with matched cell counts (Supplementary Data 3 (`completed-analysis review forms`)).

The four case studies below develop questions specific to each dataset. The spatial and perturbation analyses each drew on three method papers with distinct roles in the resulting formulation (Supplementary Note S1.5). Case 1 was identified as T1, and Cases 2–4 were identified as T2. Before execution, a search for studies related to each proposed analysis returned INSUFFICIENT, N1, N1 and INSUFFICIENT in case order. N1 denotes related methods or analytical ideas with a different question, representation or analysis. INSUFFICIENT means that the search did not resolve the plan’s relation to prior work (Method 5.1).

#### Case study 1 | Dose-resolved analysis reveals cross-dose structure in PjDHFR resistance

The deep-mutational scan of *Pneumocystis jirovecii* dihydrofolate reductase (PjDHFR) quantified variant effects as coefficients of selection (CoS) in dimethyl sulfoxide (DMSO) and two methotrexate (MTX) conditions: 8.8 *µ*M (MTX4) and 44 *µ*M (MTX20) [21]. The source study also reported a structural score, ΔΔ*G*_MTX_, derived from an MTX-bound protein model. BioMINTREL incorporated these values from the analysis files accompanying the source study’s figures. It organized these measurements as a dose-resolved genotype–phenotype landscape. MTX4- and MTX20-versus-DMSO contrasts defined four classes among 3,762 single-amino-acid substitutions: 619 cross-dose resistant, 102 high-dose-specific, 1,908 drug-sensitive and 1,133 neutral (Fig. 3a; Supplementary Note S5).

**Figure 3:**
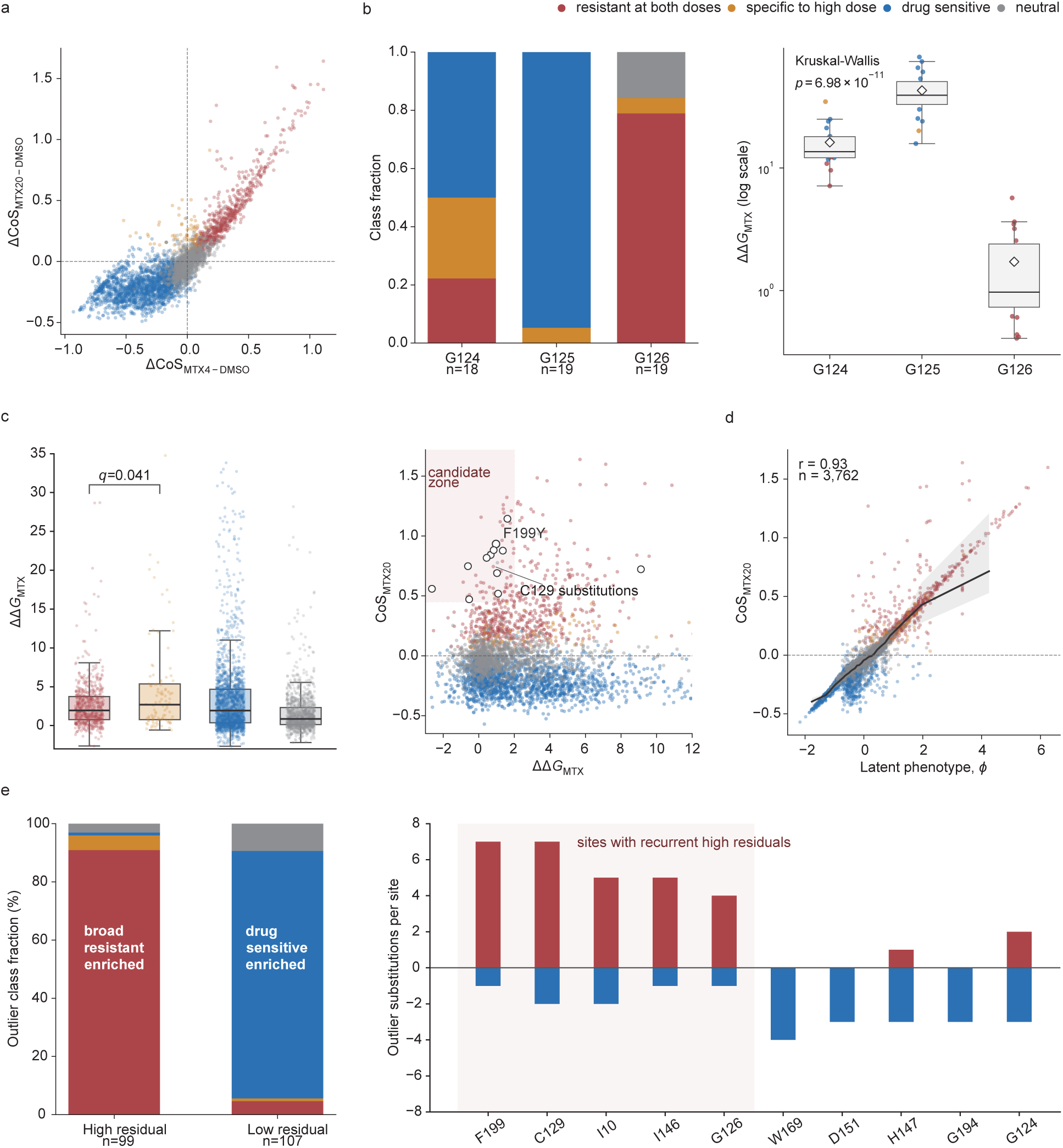
Dose-resolved analysis reveals cross-dose structure in PjDHFR resistance. **a**, Cross-dose classes defined from source-reported variant-effect values in MTX4 and MTX20 relative to DMSO. **b**, Class composition and ΔΔ*G*_MTX_ at G124–G126 (*n* = 18, 19, 19; Kruskal–Wallis *P* = 6.98 × 10*^−^*^11^). **c**, Structural-score distributions and MTX20 responses across cross-dose classes. The cross-dose-resistant versus high-dose-specific comparison has Benjamini–Hochberg-adjusted *q* = 0.041 across six class pairs. **d**, Fitted latent phenotype and MTX20 response across 3,762 substitutions. **e**, Class composition of 99 high-residual and 107 low-residual substitutions and recurrent residual positions.

Cross-dose classification resolved a pronounced local asymmetry at the adjacent G124–G126 glycine triplet. G124 showed a mixed phenotype enriched for high-dose-specific substitutions (5/18; adjusted *P* value *q* = 0.019), G125 was predominantly drug-sensitive (18/19; *q* = 4.02 × 10*^−^*^4^), and G126 was predominantly resistant across doses (15/19; *q* = 6.49×10*^−^*^8^). Mean structural scores were 16.19, 42.92 and 1.72, respectively (Kruskal–Wallis *P* = 6.98 × 10*^−^*^11^; Fig. 3b). The source study also found resistance-associated substitutions at G124 and G126 and strong constraint at G125 [21]. This agreement supports the biological relevance of the local pattern, and the dose-resolved classes show how these neighbouring residues differ across MTX conditions (Supplementary Note S5).

Cross-dose phenotypes also occupied distinct regions of the source-derived structural score, ΔΔ*G*_MTX_. Mean scores were 1.66 for neutral, 2.62 for cross-dose-resistant, 3.94 for drug-sensitive and 4.51 for high-dose-specific substitutions (Kruskal–Wallis *P* = 1.42×10*^−^*^36^). Cross-dose-resistant and high-dose-specific score distributions also differed (*q* = 0.041 across six pairwise comparisons; Fig. 3c). Cross-dose resistance was also observed at low structural scores, including F199Y (ΔΔ*G*_MTX_ = 0.97) and C129V, C129E and C129T (all below 0.84). The source study validated resistance at F199 and proposed that this distant site alters drug binding [21]. The C129 substitutions extend this low-score candidate set. Studies of bacterial DHFR link resistance to inhibitor binding, catalytic efficiency, folding stability and intracellular abundance [22]. These findings support direct measurements of these properties in the C129 variants and F199Y to explain their resistance at low structural scores (Supplementary Note S5).

Residual analysis provided a complementary view of substitutions that departed from the dominant MTX20 sequence–response trend. BioMINTREL fitted an additive MAVE-NN model, which learns genotype–phenotype maps from multiplexed variant-effect assays [23]. Its sequence-based phenotype score *ϕ* correlated with MTX20 CoS across the 3,762 substitutions (Pearson *r* = 0.929; Spearman *ρ* = 0.918; Fig. 3d). Residuals measured the difference between observed MTX20 CoS and values expected from an empirical *ϕ*–CoS calibration. Of 99 high-residual substitutions, 90 were resistant across doses. Of 107 low-residual substitutions, 91 were drug-sensitive. High residuals recurred at F199, C129, I10, I146 and G126 (Fig. 3e). The source study independently highlighted F199, I10 and the G124–G126 region [21]. Recovering these positions supports the biological relevance of the residual ranking. The dose classes, structural scores and residuals together identify C129 and I146 as candidates for testing how mutations produce resistance across MTX conditions (Supplementary Note S5).

#### Case study 2 | Flow representation resolves local constraints on spatial communication

BioMINTREL analysed five ligand–receptor pairs in a 1,657-spot breast-cancer Visium section [24]. It represented ligand-associated expression as sender supply, receptor-associated expression as local receiving capacity and spatial neighbourhoods as possible connections. Minimum-cost flow minimized total cost while respecting these capacities, allocating supply to nearby receptor-positive spots and leaving the rest unmatched [25, 26]. Comparisons used COMMOT’s spot-level scores for the same ligand–receptor pairs and CellChat’s pathway-level scores [27, 28]. Allocated sender signal closely tracked COMMOT for SPP1–CD44, CXCL12–CXCR4 and TGFB1–TGFBR1 (Pearson 0.887–0.997; Spearman 0.904–0.999), and VEGFA–KDR showed strong receiver-side concordance (Pearson 0.989; Spearman 0.997; Supplementary Note S6). CellChat supported the corresponding interactions at the pathway level. The flow representation thus retained patterns identified by established methods and added information about receiving capacity, receptor coverage and the concentration of incoming connections (Fig. 4a).

**Figure 4:**
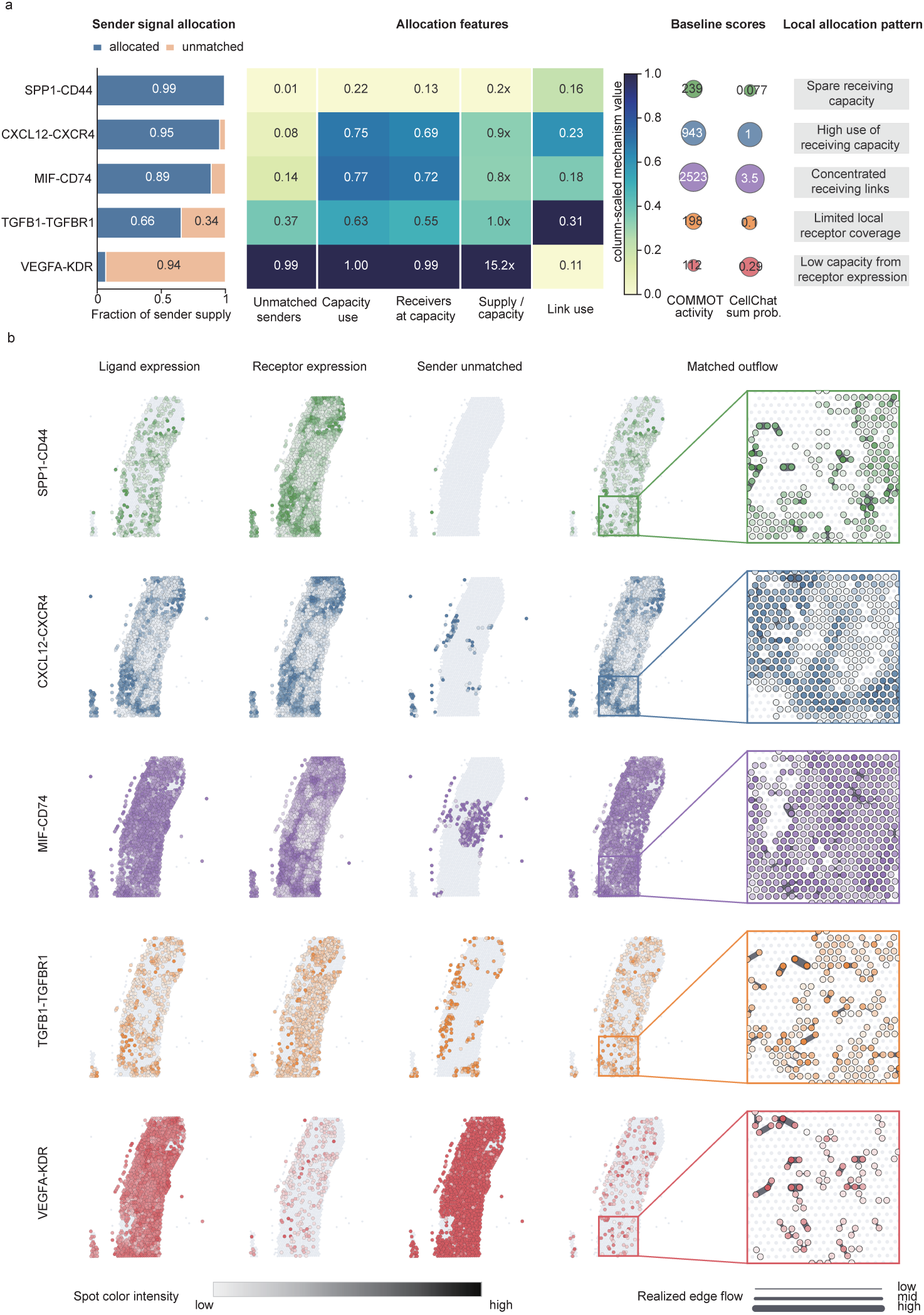
Flow representation resolves distinct local constraints on spatial ligand–receptor communication. **a**, Allocation and capacity summaries for five ligand–receptor pairs. Stacked bars show allocated and unmatched fractions of normalized sender supply. The heatmap reports the fraction of ligand-positive senders retaining unmatched signal, mean receiving-capacity use, the fraction of receptor-positive spots reaching model capacity, normalized supply/capacity and allocated/candidate incoming-link ratios. COMMOT and CellChat provide spot- and pathway-level reference scores. **b**, Ligand expression, receptor expression, unmatched sender signal and allocated outflow in the same section. The boxed regions and magnified views use the same spatial window for all five pairs. The magnified views show the 45 strongest local realized edges by flow for each pair. Within each pair, edge width scales with realized flow among the displayed edges, whereas node colour represents local realized node flow calculated from all realized edges within the common window.

SPP1–CD44 and CXCL12–CXCR4 showed similarly high allocation with different use of receiving capacity. SPP1–CD44 allocated 99.2% of sender supply, with mean receiving-capacity use of 0.219. CXCL12–CXCR4 allocated 95.5%, with 867 of 1,264 receptor-positive spots reaching model capacity and mean use of 0.750. The shared spatial window places these allocation patterns at the same tissue locations for direct comparison (Fig. 4b). These results distinguish spare capacity from high local capacity use. Previous breast-cancer studies link CXCL12–CXCR4 to migration towards ligand-rich sites and SPP1–CD44 to close macrophage–tumour contacts in chemoresistant disease [29, 30]. These findings support testing whether receptor abundance and sender–receiver proximity explain the distinct capacity-use patterns. Spatial measurements and receptor perturbation provide a direct way to examine this question (Supplementary Note S6).

MIF–CD74 showed high allocation concentrated in fewer receiving connections. The model allocated 88.9% of supply through 18.4% of candidate incoming connections. The 168 spots with high incoming allocation had higher CD74, LYZ and CD68 expression than other spots, placing this pattern in an immune-rich tissue context [31]. In mouse models of triple-negative breast cancer, tumour-derived MIF activates CD74 on dendritic cells, and CD74 blockade alters the dendritic-cell compartment and tumour growth [32]. This provides a biological basis for asking whether CD74-positive dendritic cells account for the concentrated incoming signal. Mapping dendritic-cell localization and CD74 activity within these spots, followed by CD74 perturbation, would test this interpretation (Supplementary Note S6).

TGFB1–TGFBR1 and VEGFA–KDR separated two forms of limited allocation. TGFB1–TGFBR1 allocated 65.5% of supply at a nearly balanced normalized supply-to-capacity ratio of 1.048. Senders retaining unmatched signal had fewer receptor-positive neighbours than fully allocated senders (2.65 versus 5.39) and a greater nearest-receptor distance (306.9 versus 153.0 spatial units), consistent with limited local receptor coverage. Previous breast-cancer analyses localized TGFB1–TGFBR1/TGFBR2 activity to tumour–stroma interfaces [33]. This supports examining whether receptor coverage across these boundaries explains incomplete allocation despite balanced total supply and capacity. VEGFA–KDR had a normalized supply-to-capacity ratio of 15.222: 6.5% of supply was allocated and 291 of 294 receptor-positive spots reached model capacity. Its 34 spots with high incoming allocation expressed higher KDR and vascular markers, including PECAM1 and PLVAP [34]. Previous studies reported VEGFA–KDR coexpression in breast tumours and reduced angiogenesis after VEGF–KDR blockade [35, 36]. Together with the vascular markers, this evidence supports examining whether receptor availability in vascular regions limits local VEGFA allocation. Histological mapping and TGFBR1 perturbation could test whether local receptor coverage explains the TGFB1 pattern. Measurements of KDR abundance and vascular response, followed by KDR perturbation, could test the VEGFA interpretation (Supplementary Note S6). Across the five pairs, the model resolves spare capacity, high capacity use, concentrated receiving connections, limited receptor coverage and low receptor-derived capacity. These patterns define targets for colocalization, receptor perturbation and replication across tissue sections.

In the shared window, MIF–CD74 allocation is widespread and VEGFA–KDR allocation forms smaller local groups (Fig. 4b). The whole-section summaries relate these patterns to concentrated receiving links and low receptor-derived capacity, respectively.

#### Case study 3 | State-graph analysis resolves shared and alternative perturbation routes

The source Perturb-seq study measured single-cell transcriptional responses to genetic perturbations of the unfolded protein response (UPR) [37]. BioMINTREL represented the 11 expression-defined states (C0–C10) as a graph and applied transition-path theory (TPT) to compare routes across 15 perturbations [38–40] (Fig. 5a). Connections reflected similarities between the cells’ expression profiles. BioMINTREL calculated UPR scores using gene sets drawn from the source paper’s figures and text to summarize the overall response and its branches. For each perturbation, higher control-cell fractions and lower UPR scores favoured starting states. Higher fractions of cells carrying that perturbation and higher UPR scores favoured target states. BioMINTREL weighted connections to favour movement towards states with a higher fraction of cells carrying that perturbation and stronger UPR activity. TPT assigns each state a forward committor, the probability of reaching the target before returning to the source. Reactive flux measures each directed connection’s contribution to paths between these sets. BioMINTREL used the incoming and outgoing net flux to score intermediate states (Supplementary Note S7).

**Figure 5:**
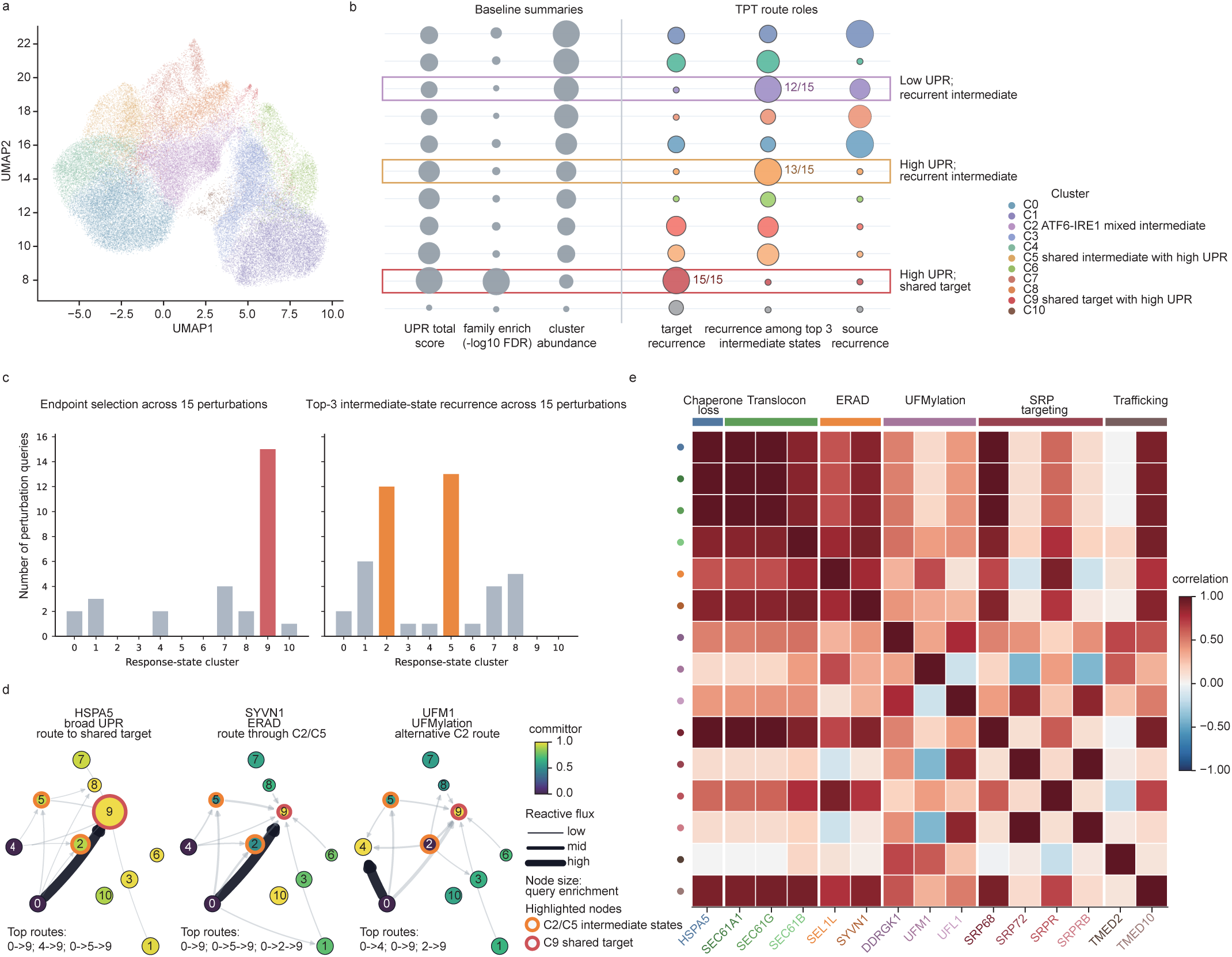
State-graph analysis resolves shared and alternative Perturb-seq routes. **a**, UMAP[51] of the 11 transcriptional response states. **b**, Baseline state summaries alongside target, intermediate-state and source recurrence in the TPT representation. **c**, C9 enters the target set in 15/15 queries. C5 and C2 occur among the leading intermediate states in 13/15 and 12/15 queries. Procedure-matched permutation controls are reported separately. **d**, Representative HSPA5, SYVN1 and UFM1 routes; node colour denotes the forward committor, node size follows the query-enrichment scaling shown in the panel and edge width denotes net reactive flux. **e**, Pearson correlations of 11-state committor vectors across perturbation queries.

C9 contained 1,261 cells, had the highest mean aggregate UPR score (2.259) and entered the target set in all 15 perturbations. Chaperone-loss perturbations were enriched 5.55-fold in C9 (odds ratio 17.47), with HSPA5 accounting for 33.1% of its cells. C5 and C2 appeared among the three highest-scoring intermediate states in 13/15 and 12/15 perturbations, respectively (Fig. 5b,c). C5 occupied a recurrent position near the high-UPR target region. C2 had a low aggregate UPR score (−0.227) and participated frequently in inferred routes. Its prominence therefore came from its position along these routes. The graph adds information about connections between response states beyond a ranking by UPR intensity.

The starting states, target states and connection directions depend on how each perturbation is distributed across the graph. We therefore asked how much of the recurrence of C9, C5 and C2 could arise from these choices and the shared graph. A permutation analysis repeated the full path-definition procedure 5,000 times, permuting labels among cells from the 15 selected perturbations, retaining each perturbation’s cell count (Supplementary Note S7). Their observed recurrence fell within the resulting control distributions. C9, C5 and C2 thus provide common reference states for comparing inferred routes. Time-resolved or independent measurements can test whether cells also pass through these states during a response.

Route profiles grouped biologically related perturbations (Fig. 5d,e). The dominant C0→C9 route appeared in 9/15 perturbations. SEL1L and SYVN1, which act in the same endoplasmic-reticulum-associated degradation (ERAD) machinery [41], shared this route and C5 as a leading intermediate state (committor correlation 0.805; edge-flux cosine similarity 0.839). HSPA5 and SEC61A1 had nearly identical graph profiles (committor correlation 1.000; flux similarity 0.999), with SEC61G nearby. SEC61B shared the C0→C9 route with secondary routes through C2 and C5. HSPA5 and the Sec61 translocon both regulate endoplasmic-reticulum stress sensors [42, 43], supporting this functional grouping. UFMylation, the attachment of UFM1 to target proteins, supports translocation-associated ribosome quality control and endoplasmic-reticulum homeostasis [44]. Its perturbations followed distinct routes: UFL1 followed C4→C0→C9, UFM1 followed C0→C4 and DDRGK1 followed C0→C9. These established functions support the biological basis of the route groupings (Supplementary Note S7). The distinct inferred routes within each pathway identify states at which time-resolved measurements can test how the responses diverge.

#### Case study 4 | Upper-tail analysis reveals sampling requirements for rare-cell prediction

The Watermelon lineage-tracing data link early cell states to later clonal outcomes across thousands of PC9 lung-cancer lineages [45]. Gene-programme scores summarize expression of defined gene sets. For each programme, BioMINTREL used an empirical upper-tail summary: the mean of the highest 5% of scores within each lineage. This summary was motivated by conditional value at risk (CVaR), a statistic that summarizes a specified distributional tail [46–48]. This representation tests whether rare high-state cells provide information about later lineage size beyond early abundance and average state. BioMINTREL selected xenobiotic metabolism, glutathione metabolism and reactive oxygen species (ROS) defence based on prior evidence on rare states and drug resistance [49, 50] and pathway findings in Watermelon’s main and Extended Data figures. Notch and interferon-alpha served as biological controls [45] (Supplementary Note S8).

Day-3 sampling strongly constrained the upper-tail statistic. The dataset contained 11,348 day-3 cells across 5,177 lineages; 64.0% of lineages had one observation, 90.0% had fewer than five and 97.3% had fewer than ten (Fig. 6a). With at most 20 observations per lineage, the highest 5% contains only the highest-scoring cell. The empirical top-5% mean therefore requires at least 21 observations to include more than one cell, a condition met by 23 lineages. Across the seven scores used to assess sampling depth (the five programmes, fatty acid metabolism and the cell-cycle score), the top-5% mean equalled the lineage maximum in 99.58% of summaries (Fig. 6b). Measuring a tail formed by several cells therefore requires deeper sampling within each lineage.

**Figure 6:**
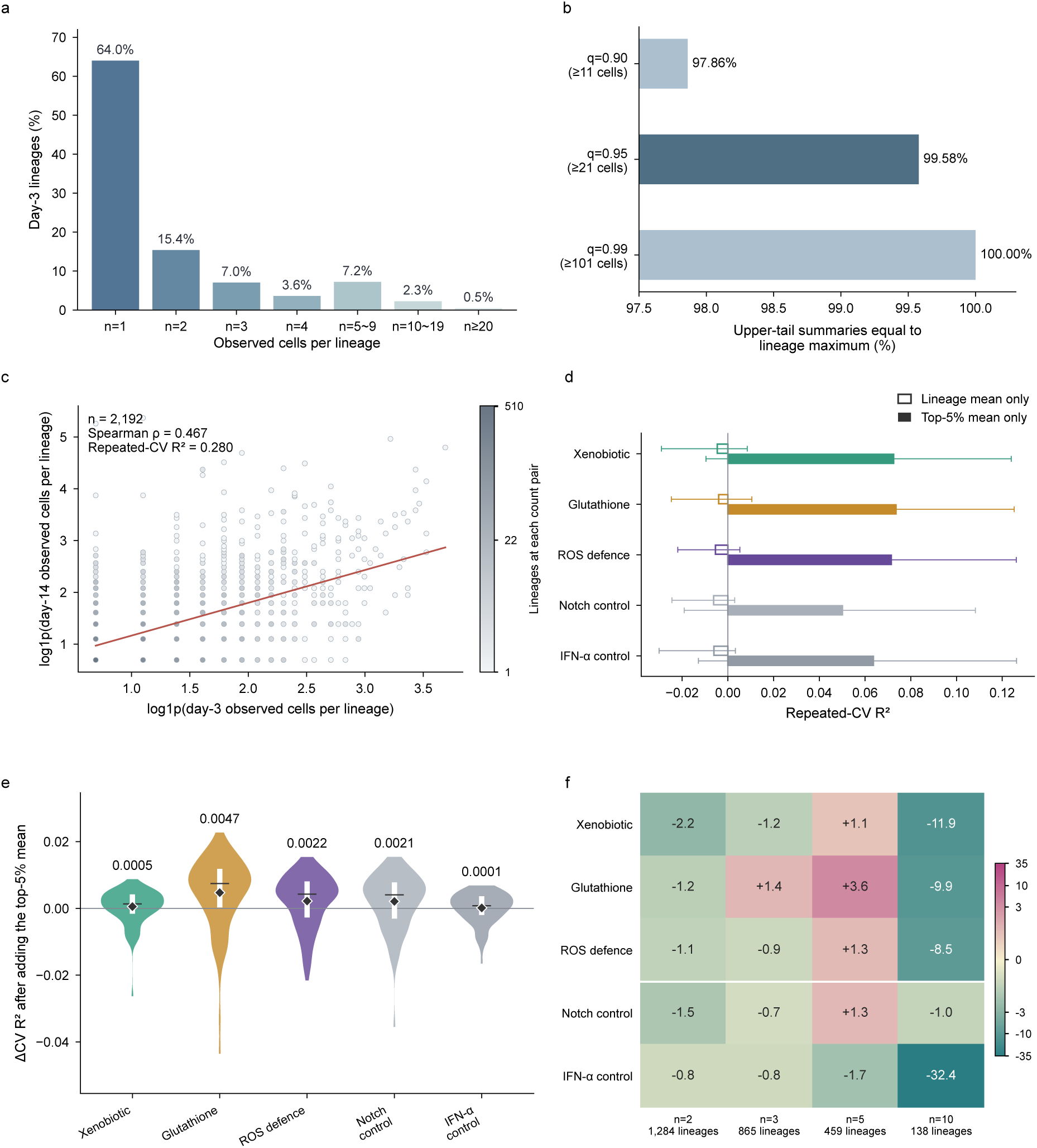
Upper-tail analysis reveals the sampling requirements for rare-cell prediction. **a**, Observed day-3 cells per lineage. **b**, Fractions of lineage–programme summaries in which the empirical upper-tail mean equals the maximum. **c**, Early observed lineage abundance and later lineage size (Spearman *ρ* = 0.467). **d**, Mean-only and nominal top-5%-mean-only prediction of later lineage size for candidate and control gene programmes. Bars show means, and whiskers span the central 95% of 100 cross-validation fold scores. **e**, Paired fold-wise change in cross-validated *R*^2^ after adding the top-5% summary to abundance and lineage mean. Violins show 100 matched fold differences per programme; diamonds mark means, white segments show interquartile ranges and black lines mark medians. **f**, Equal-cell resampling compares mean-plus-tail with mean-only prediction at matched within-lineage cell counts. Cell annotations show mean increments in units of 10*^−^*^3^.

Early lineage abundance was associated with later size (Spearman *ρ* = 0.467; mean repeated-cross-validation *R*^2^ = 0.280; Fig. 6c). Larger early samples also have more chances to contain a high-scoring cell. BioMINTREL therefore assessed the tail after accounting for abundance and the lineage mean. Among the 2,192 lineages observed at both day 3 and day 14, adding the top-5% mean to abundance and lineage mean increased mean fold-wise cross-validated *R*^2^ by 0.0005, 0.0047 and 0.0022 for xenobiotic metabolism, glutathione metabolism and ROS defence, and by 0.0021 and 0.0001 for the controls (Fig. 6e). These comparisons mainly assess the highest-scoring sampled cell because the top-5% mean usually contains one cell. Comparisons at matched cell counts further tested what the highest-scoring cell adds beyond the lineage mean (Fig. 6f; Supplementary Note S8). A direct test of the proposed upper-tail contribution therefore needs enough early observations within each lineage to include several high-state cells.

The 21-observation threshold marks the first point at which the chosen top-5% statistic can include multiple cells. Stable estimation requires deeper and more even early lineage sampling together with independent biological replication. At matched sampling depth, this design can test whether high-state cells provide reproducible information about later outgrowth beyond lineage abundance and average state. BioMINTREL thus uses existing data to define both a biological question and the sampling needed to answer it.

## Discussion

BioMINTREL expands the questions that researchers can ask of existing biological data. The same measurements can be organized as a matrix, graph or distribution, each exposing different relationships. BioMINTREL searches for methods from other disciplines that use these representations and defines how their computational requirements can be met by biological measurements. It can use relationships among datasets to form joint questions and combine ideas from several methods to develop analyses. This opens analytical directions beyond those considered in the original study.

The central step is method–data correspondence. Similarity between scientific fields can suggest a candidate method, but a useful transfer requires the data to provide meaningful inputs and relationships, the core calculation to retain its role, and the output to address a biological question. Information from paper text, figures, tables and supplementary materials helps define these inputs and their biological meaning. Defining these elements turns a possible analogy into an analysis that can be formulated and tested. It also explains why analytical structure complements scientific context when searching for methods: a method developed for a different topic can still require the mathematical objects and relationships available in biological data. Scientific criteria and complete analysis plans then determine which possibilities can support a concrete study.

The four case studies show how a new representation can change the questions asked of existing data. A dose-resolved genotype–phenotype representation distinguishes mutation effects that persist across drug concentrations from those that emerge under stronger selection. Spatial allocation asks how local supply, receptor capacity and accessibility constrain possible communication, while a state graph compares shared endpoints, recurrent intermediate states and alternative routes through perturbation responses. The lineage case shows another role for a new question: it can identify the measurements needed to answer it. With the available early sampling, the upper-tail mean usually represents a single observed cell. Deeper within-lineage sampling is therefore needed to test whether rare high-state cells provide information beyond abundance and average state. Together, these cases show how new analyses can reveal biological relationships and direct subsequent research toward the evidence needed to test them.

The evaluation supports the framework from several angles because useful method transfers cannot be captured by a single fixed reference. Across two language models, explicit correspondence brought judgments closer to expert consensus than direct model assessment and recovered more expert-supported pairings. Analytical-structure retrieval, scientific prioritization and independent plan generation helped bring less obvious methods into consideration and develop them into complete plans. Later literature provided a separate retrospective reference for related applications. Blinded expert reviews examined the proposed plans and completed analyses, supporting their analytical novelty and value for further investigation. Together, these findings connect the identification of plausible method applications to studies that can add biological information.

BioMDC-Bench fixes tasks and candidate methods so that expert judgments, direct LLM judgments and BioMINTREL can be compared on the same combinations. Extending this evaluation to other biological areas, datasets and method families, including questions formed from multiple sources, would test how widely BioMINTREL can expand the research directions considered. The latter literature analysis is retrospective and may include studies represented in model training, so we interpret it as consistency with research activity. Prospective studies can formulate questions and analysis plans before related findings appear, then test them using independent data and targeted experiments. This would also examine promising method–data combinations without close published precedents, since published work reflects the questions and approaches researchers previously chose to pursue.

By defining the question, representation, core computation, controls and required evidence before an analysis, BioMINTREL shows what an existing dataset can support and what further measurements would make a question testable. This matters for accumulated datasets whose original analyses addressed only part of the information they contain. Methods developed in other fields can reveal additional relationships, and their requirements can clarify which comparisons or experiments should come next. BioMINTREL thus connects existing biological data to new questions, analyses and focused follow-up studies.

## Methods

BioMINTREL develops new biological analyses by linking existing biological data with computational methods from other fields. It starts from one or more data papers and one or more method papers, with an optional biological question and method name. If no biological question is specified, BioMINTREL can identify a question supported by the available data. It then ranks candidate methods, determines how each method can be applied to the data, and develops supported method–data pairs into complete analysis plans. Following selected plans, BioMINTREL will execute the analysis, then combine the resulting measurements, comparisons and relevant literature into biological interpretations. Algorithm 1 summarizes the procedure.

### 1 Inputs

BioMINTREL takes one or more data papers and one or more method papers as inputs. In the notation below, *B_i_* denotes the data papers considered together for a task, and *A_j_* denotes the papers supporting a candidate method *m_j_*, which may combine ideas or computations from several methods. Data papers describe the biological samples, experimental design, measured variables and their biological meaning. We denote the measurements and sample annotations selected for the task by *d_i_*, which may draw on several datasets. Method papers describe input types, core operators, required data relations and computational outputs. BioMINTREL examines whether data sources complement one another or connect biological stages. It also considers whether methods offer complementary operations, form successive steps in which one method’s output supplies another’s input, serve as comparators in a benchmark, or contribute components and ideas to a combined model (Supplementary Note S1.5).

The specific biological question 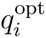 and method names *µ_j_* are optional inputs. When the user provides a biological question, BioMINTREL first checks whether the supplied data support this question, then organizes the analysis around directions supported by the data. When the user does not specify a question, BioMINTREL summarizes the available observations, relations and potential uses from the data papers, and can select a question supported by the data as the starting point for analysis. Before deciding on a specific task, the user can let the agent use the two types of papers to explore what other questions the data can answer. The method sources are denoted by 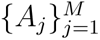, where *M* counts candidate methods and *µ_j_* is the optional name associated with *m_j_*. Each *A_j_* may contain several papers. Thus, the task–method notation refers to the resulting analytical choices, and BioMDC-Bench fixes one predefined task and one candidate method per pair for comparison. ∅ denotes an unspecified optional input. The algorithm uses retrieval weights *ω_c_, ω_a_*, a maximum of *k* methods per task, *J* initial plans per pair and at most *R* completion attempts.

BioMINTREL then connects the data with methods, establishes new data representations and identifies biological questions that the existing data can answer. It specifies which computational objects the data can provide and how to define the relations between these objects. For example, the same spatial expression data can be organized into an expression matrix or combined with spatial coordinates to construct an adjacency graph. Different data representations allow different computations, making it possible to pose different biological questions using the same data.

### 2 Extracting paper content and ranking candidate methods

#### 2.1 DatasetCard and MethodCard: Generating dataset and method cards

BioMINTREL uses the main text and information from figures, tables and available supplementary materials of the data papers in *B_i_*, extracts the biological samples, measured variables, experimental conditions, sample relations, existing annotations and potential analytical uses, and produces a dataset card *D_i_*. The dataset card *D_i_* also distinguishes directly measured information from quantities that require calculation, and lists the representations supported by the data. BioMINTREL extracts input types, required relations, core operators, outputs and applicable assumptions from the text, figures, tables and supplementary materials of the method papers in *A_j_* to produce a method card *H_j_*. For example, the inputs of a graph method require explicit definitions of nodes, connections and measurements on the nodes, while a matrix factorization method requires explicit meanings for the rows and columns of the matrix and constraints on its values. For a combined candidate, the method card distinguishes the source computations and outputs retained in the formulation and records each method’s role. Both types of cards retain the corresponding evidence from the original sources for subsequent correspondence assessment. Reported gene selections, screening results and annotations can enter the analysis as defined inputs (Supplementary Note S1.6).

#### 2.2 RankCandidateMethods: Ranking by Scientific context and Analytical structure

Candidate methods come from the supplied set of method papers. BioMINTREL searches the PubMed Central open access literature[52], locates relevant papers by their titles and abstracts, and then reads the full text and supplementary materials.

To rank candidate methods, BioMINTREL constructs two retrieval texts from the dataset card *D_i_* and method card *H_j_*: Scientific context and Analytical structure. Scientific context retains the original domain information from the papers. For the data, it includes biological samples, measurement types, experimental conditions and research questions. For a method, it includes the problem originally addressed by the method and its main applications. Therefore, Scientific context mainly reflects whether the data and method are close in their original application domains. Analytical structure removes specific domain and application names and retains only computation-related features. For the data, it describes the mathematical objects that the data can form, such as matrices or graphs, as well as observational units, existing relations, structural constraints and targets that can be analyzed. For a method, it describes the mathematical objects on which the core operator acts, the constraints it requires and the types of results it produces. For example, non-negative matrix factorization can be represented as “non-negative matrix—matrix factorization—factor output”, without retaining its original application context in image or text analysis[53].

##### Algorithm 1 BioMINTREL analysis from data and method papers

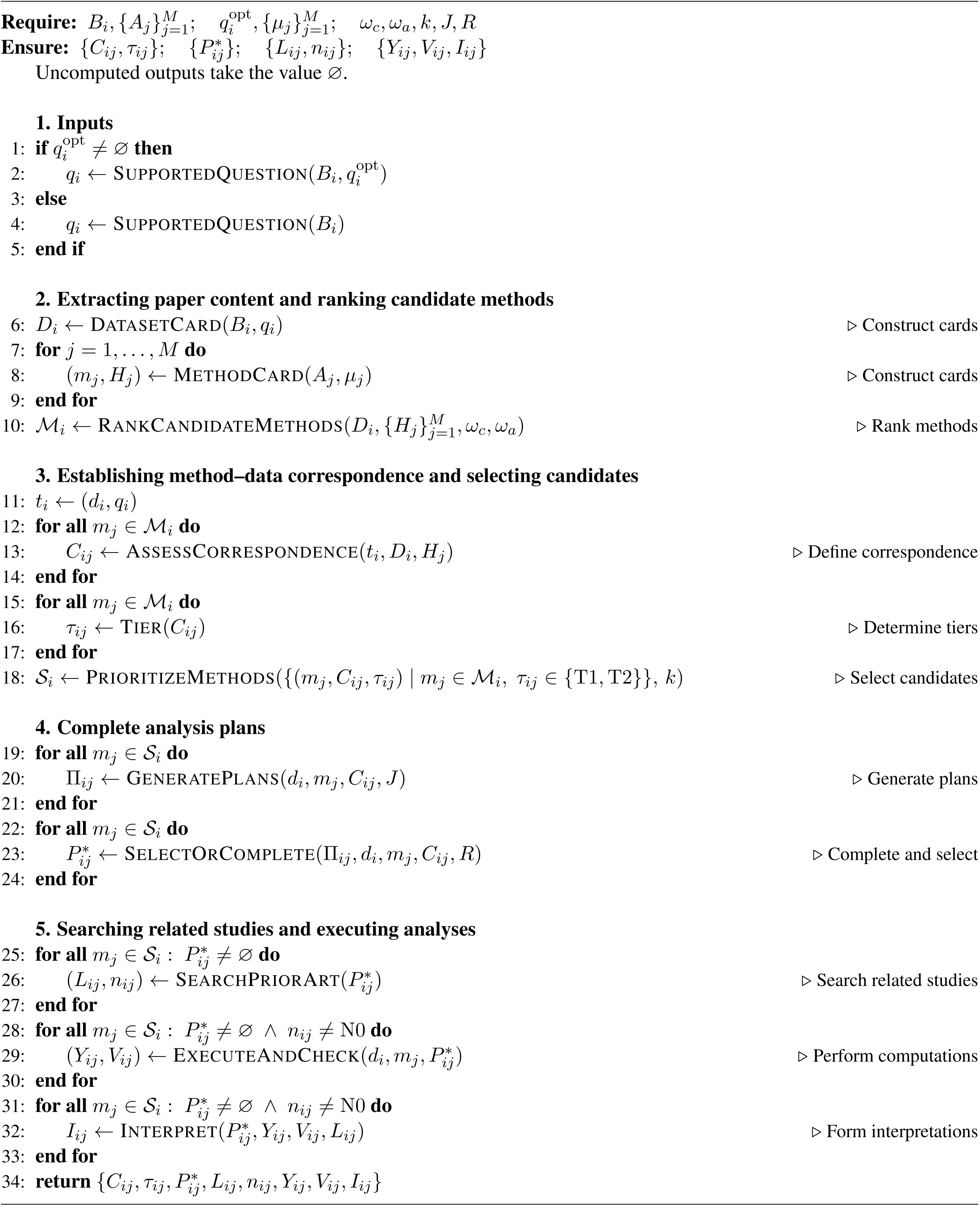

Denote the data used in the current analysis by *d_i_*, the starting biological question by *q_i_*, and the *j*th candidate method by *m_j_*. BioMINTREL uses SPECTER2, a scientific text encoder [54], to encode the two retrieval texts separately and calculates the cosine similarity between the corresponding data-side and method-side texts. For *d_i_* and *m_j_*, the Scientific context similarity and Analytical structure similarity are denoted by 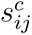 and 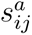, respectively. The two weights are denoted by *ω_c_* and *ω_a_*, respectively, and the combined retrieval score is denoted by *r_ij_*:

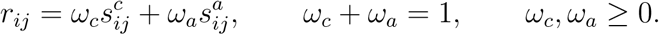

The Scientific context similarity 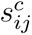 reflects the closeness of the original applications, and the Analytical structure similarity 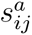 reflects the closeness of the computational features. RankCandidateMethods uses *r_ij_* to rank candidate methods *m_j_* from highest to lowest. The resulting ordered method list is denoted by 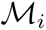.

### 3 Establishing method–data correspondence and selecting candidates

#### 3.1 AssessCorrespondence: Defining inputs, relations and the biological question

The data *d_i_* and starting question *q_i_* form the task *t_i_* = (*d_i_, q_i_*). BioMINTREL pairs each task *t_i_* with a candidate method *m_j_* to form a task–method pair (*t_i_, m_j_*).

The set of input objects that method *m_j_* requires before its core computation is denoted by *U_j_*, specifying which data objects the method requires. Let *Ɛ_j_* denote the set of data relations that the method requires between these input objects, specifying which relations these objects need to have. For example, for a graph method, nodes and measurements on the nodes can belong to *U_j_*, and adjacency relations between nodes belong to *Ɛ_j_*. The core computation of the source method is denoted by *K_j_*, including its key operations, objective function and constraints. The output originally defined by the method is denoted by *o_j_*. The mathematical constraints of the method are first recorded as part of *K_j_*. If a constraint also requires the target data to provide specific inputs or data relations, the corresponding requirements are additionally included in *U_j_* or *Ɛ_j_*.

BioMINTREL determines how *U_j_* and *Ɛ_j_* are provided by the data *d_i_* item by item, while specifying how *K_j_* acts on these inputs, what biological meaning *o_j_* has and what evidence can test this result. This Method–data correspondence is denoted by *C_ij_*, and the final biological question determined through correspondence assessment is denoted by *q_ij_*. Thus, *C_ij_* specifies how *d_i_* meets the requirements in *U_j_* and *Ɛ_j_*, and how the resulting computation supports *K_j_* and *q_ij_*.

*C_ij_* needs to be specified in terms of actual computation. Each item in *U_j_* and *Ɛ_j_* requires a clear data source or computational rule. For example, a spatial graph method requires nodes, measurements on the nodes and connections between nodes. Taking spatial transcriptomic data with single-cell coordinates as an example, cells can serve as nodes, expression levels as measurements on the nodes, and spatial coordinates provide the cell positions. However, this information does not yet directly determine the graph structure. BioMINTREL also needs to specify which cells are connected and how the connections are calculated from spatial positions or other observed quantities. If *Ɛ_j_* lacks a clear basis in the data or computational rules, even if a conceptual analogy is possible, this correspondence in *C_ij_* remains incomplete.

Number the items in *U_j_* ∪ *Ɛ_j_* in sequence, and denote the construction status of the *h*th requirement by *σ_ijh_*. **Direct observation (observed)** means that the required information already exists in the original measurements or sample annotations of *d_i_*. **Fixed construction (deterministic)** means that the required information can be uniquely obtained from *d_i_* according to a predetermined rule. For example, once the distance definition is specified, intercellular distances can be calculated directly from spatial coordinates.

*U_j_* or *E_j_* sometimes has multiple reasonable definitions. In this case, BioMINTREL can use a **bounded choice (bounded)**: specify the range of candidates in advance and define a selection criterion that is independent of the final analysis results. For example, several neighborhood scales can be specified in advance, and one can then be selected according to predetermined graph structure requirements. A bounded choice requires an explicit candidate set and selection rule. The subsequent plan determines a specific value on this basis and checks whether the value belongs to the candidate set specified in advance. The selection rule cannot depend on which setting produces more significant results or results that better match expectations.

If the *h*th requirement needs a new definition without a basis in the data, an analysis choice within an open-ended range, or another as-yet undefined analytical problem to be solved first, BioMINTREL records it as **missing/open**. See Supplementary Note S1.1 for the specific decision principles and examples of category boundaries.

Thus, *σ_ijh_* describes the construction status of a single requirement in *U_j_* or *Ɛ_j_*, while *C_ij_* summarizes all correspondences for the entire task–method pair. A *C_ij_* can contain observed, deterministic and bounded items at the same time. If it still contains a missing/open item that prevents *K_j_* from being executed, the overall correspondence still has a critical gap.

In addition to the item-level correspondences for *U_j_* and *Ɛ_j_*, *C_ij_* records whether *K_j_* is retained after transfer, the relation between the final question *q_ij_* and the starting question *q_i_*, and the adjustments required during application. If *q_i_* remains unchanged during correspondence assessment, then *q_ij_* = *q_i_*. The data representation determined in *C_ij_* for use in subsequent plans is denoted by *X_ij_*. *C_ij_* also specifies how *o_j_* addresses *q_ij_* and which comparison tests that output.

For the core computation *K_j_*, BioMINTREL determines whether the analysis after transfer still retains its key operations, objective function and constraints. Parameter settings within a defined range can be used within *m_j_*, as long as they do not change *K_j_*. If another method has replaced the main computation, or the continued use of the key operations of *K_j_* cannot be confirmed, *K_j_* is not considered retained. Method 3.3 further checks whether *K_j_* actually provides key results for the main analysis that answers *q_ij_*. See Supplementary Note S1.2 for the calculation criteria and Supplementary Note S1.4 for the scientific-priority example.

The main result used to answer *q_ij_* is denoted by *y_ij_*. *y_ij_* can use the output *o_j_* originally defined by the method, obtain an equivalent result through a fixed transformation, or add a new interpretation specified in advance. If *o_j_* only provides a new data representation, such as an embedding, further explanation is needed of how this representation is used to answer *q_ij_*. This information is recorded with *C_ij_* and checked further during subsequent candidate selection and plan generation. See Supplementary Note S1.2 for specific definitions.

Finally, BioMINTREL specifies a comparison appropriate to *y_ij_*, with the observations or comparison objects denoted by *v_ij_*. For example, for a model that predicts responses across multiple samples, samples not involved in model development can be designated for validation in advance. Final held-out observations must remain separate from model fitting and selection[55]. Descriptive comparisons and predefined null or sensitivity analyses test the stated properties of a representation. See Supplementary Note S1.4 for the comparison criteria. The correspondences for *U_j_* and *Ɛ_j_*, the retention of *K_j_*, the link between *q_ij_* and *q_i_*, and the overall form of adaptation determine the tier in the next section. The interpretation of *y_ij_* and the validation approach are recorded with *C_ij_* for subsequent candidate selection and plan completion. Specific data splits, analysis units and computational steps are further specified during subsequent plan generation.

#### 3.2 Tier: Determining adaptation tiers

Based on the item-by-item correspondences in *C_ij_*, BioMINTREL determines an adaptation tier for (*t_i_, m_j_*), denoted by *τ_ij_* ∈ {T1, T2, T3}, to describe the overall form of method transfer.

T1: Direct application. All requirements in *U_j_* and *Ɛ_j_* have explicit correspondences, and each *σ_ijh_* is observed or deterministic. *q_i_* remains unchanged, *K_j_* is retained, and the overall application requires no additional adjustments or only minor adjustments. See Supplementary Note S1.3 for the specific conditions.

T1 allows minor adjustments that do not change *q_i_* or *K_j_*, such as changes in matrix orientation, fixed unit conversions and other minor adjustments. *U_j_* and *Ɛ_j_* can also be constructed through fixed rules, but the meanings of *q_i_*, *K_j_* and E*_j_* remain unchanged. See Supplementary Note S1.3 for the specific boundaries.

For example, *m_j_* requires a non-negative “sample × feature” matrix and extracts shared patterns through matrix factorization. If *d_i_* already has this matrix form and *q_i_* is precisely to identify shared expression patterns, then samples, genes and expression values can correspond directly to the inputs in *U_j_*, and *m_j_* can be executed without establishing new data relations.

T2: Constrained cross-disciplinary application. All requirements in *U_j_* and *Ɛ_j_* have established correspondences, *K_j_* is retained, and *q_ij_* still has a clear link to *q_i_*, but the overall application goes beyond the direct correspondence of T1. Typical situations include a bounded *σ_ijh_*, the need to reformulate *q_i_* as a clearly related *q_ij_*, or other adjustments with defined boundaries required to apply *m_j_*. See Supplementary Note S1.3 for the specific conditions.

The scope and principles of these changes need to be specified before analysis. If multiple candidate constructions are involved, the range of candidates and a selection rule independent of the final results also need to be specified in advance. They cannot rely on open-ended assumptions, and the definition to use cannot be chosen after seeing the final results. See Supplementary Note S1.3 for the relation between the local bounded construction status *σ_ijh_* and the overall tier *τ_ij_*.

For example, a graph method requires nodes, node attributes and adjacency relations. Spatial transcriptomic data can directly provide cells, expression levels and spatial coordinates, but adjacency relations need to be established further. If the candidate neighborhood scales are specified in advance and the specific scale is chosen according to a principle independent of the final biological conclusion, then all graph inputs can be explicitly constructed. The adjacency relation in E*_j_* in this example requires selection among multiple candidate constructions according to a predetermined rule, so the corresponding *σ_ijh_* = bounded. When all other adaptation conditions are satisfied, the entire pair (*t_i_, m_j_*) has *τ_ij_* = T2.

T3: Required correspondence not yet established. When at least one requirement in *U_j_* or *Ɛ_j_* cannot be met through observed, deterministic or bounded construction, *K_j_* cannot be retained, *q_ij_* loses its clear link to *q_i_*, or the application of *m_j_* still requires open-ended definitions of new data relations, questions or other key content, the pair (*t_i_, m_j_*) has *τ_ij_* = T3.

See Supplementary Note S1.3 for the full conditions defining T1, T2 and T3. T1 and T2 distinguish the changes required to apply a method and do not indicate higher or lower scientific value or adaptation quality. In T1, *m_j_* is already relatively close to the existing form of *d_i_* and to *q_i_*. T2 can reorganize the existing measurements in *d_i_*, establish data relations with a clear basis, or reformulate *q_i_* as *q_ij_*, thereby expanding the analyses that *d_i_* can support. The cross-disciplinary innovation of interest to BioMINTREL is mainly reflected in the establishment of these new data representations and questions in T2. Therefore, the classification evaluation combines T1 and T2 and compares them with T3 to assess the ability of BioMINTREL to identify whether *m_j_* and *d_i_* have established an executable correspondence *C_ij_*.

#### 3.3 PrioritizeMethods: Selecting candidates using three scientific criteria

Pairs (*t_i_, m_j_*) with *τ_ij_* of T1 or T2 can proceed to subsequent plan generation. The same task *t_i_* usually has multiple feasible methods *m_j_*, so BioMINTREL further considers three aspects to determine which *C_ij_* should be given priority for development into complete plans. This step only determines the order of candidate pairs (*t_i_, m_j_*) and does not change the previously determined *τ_ij_*.

The first criterion checks whether the correspondences in *C_ij_* established for *U_j_* and *Ɛ_j_* have a clear biological basis. Each correspondence needs to specify which measured quantity it comes from and why those measurements can serve the role required by the method. For example, when spatial distance between cells serves as an edge weight in a graph, the correspondence needs to specify that the distance is calculated from spatial coordinates and explain why this distance is suitable for representing cell relations in *q_ij_*.

The second criterion checks whether *K_j_* actually contributes to the main result *y_ij_*. *K_j_* needs to produce *y_ij_* or key intermediate results. BioMINTREL determines whether, if *K_j_* is removed, the remaining steps can already independently accomplish the same analytical objective. This criterion distinguishes actual method transfer from simply adding a method step to the analysis. See Supplementary Note S1.4 for specific examples.

The third criterion checks whether the comparison with *v_ij_* tests *y_ij_* without treating a quantity created by the adaptation as independent evidence. Final held-out observations must remain separate from model fitting and selection. For example, held-out samples can test predictions, but cannot also be used for model fitting or to select the analysis settings that produce the best predictions. See Supplementary Note S1.4 for specific examples.

BioMINTREL uses these three criteria to rank candidates with the same *τ_ij_*. Candidates with fewer identified problems receive priority for plan generation. For task *t_i_*, at most *k* methods *m_j_* are selected. The selected method list is denoted by S*_i_*.

### 4 Complete analysis plans

#### 4.1 GeneratePlans: Independently generating candidate plans

For each selected pair (*t_i_, m_j_*), where *m_j_* ∈ S*_i_*, BioMINTREL develops it into a specific analysis plan. Each pair retains the previously determined *q_ij_*, data representation *X_ij_*, core computation *K_j_*, the correspondences in *C_ij_* for *U_j_* and *Ɛ_j_*, and the main result *y_ij_* and validation target *v_ij_*. BioMINTREL uses this shared material to organize the computational steps, connects *d_i_*, *U_j_*, *K_j_* and *y_ij_*, and specifies data transformations, key parameters, data splits and comparison approaches that have not yet been determined. Each step needs to specify which data it uses, which computation it performs and how its result enters the next step. When an intermediate representation is produced, the plan also needs to specify how to obtain *y_ij_* from that representation to complete the target prediction or biological comparison.

One round of plan generation (Formulation) means producing a candidate analysis plan from the shared material described above. The plan from the *ℓ*th independent generation is denoted by 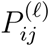. Each selected pair (*t_i_, m_j_*) independently generates *J* initial candidates, denoted by

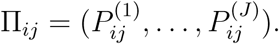

Within Π*_ij_*, each 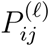 is generated using the same material, without reading the contents of the other candidates. Each 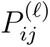 separately specifies the analysis choices that have not yet been determined, while retaining the previously determined *q_ij_*, *m_j_* and *C_ij_*.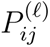 states the hypotheses to be tested, the expected findings and which observations would support or change the corresponding interpretation. See Supplementary Note S2.1 for the settings used for repeated generation.

Each 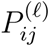 specifies the analysis unit, the object directly modelled or compared. Predictive plans also specify an independent evaluation unit: the observations that must remain together when splitting fitting and test data. For example, the analysis can use single-cell measurements and split data by lineage, so that cells from the same lineage enter the same group. These plans distinguish observations used to build the model from those used to test the results. For other comparisons, the plan identifies the observations and grouping used by the predefined comparator, null model or sensitivity calculation. See Supplementary Note S2.2 for the units and comparisons.

### 4.2 SelectOrComplete: Checking completeness, completing and selecting plans

Let *P* denote the candidate plan currently under examination. BioMINTREL compares *P* item by item with the previously determined *q_ij_*, *m_j_* and *C_ij_*. It identifies the required evidence in *P* and checks the following five requirements using uniform rules.

| Completeness requirement | Content that needs to be specified |
| --- | --- |
| **The data, question and main result are consistent** | Specify which data in $d_i$ are used, which objects are analyzed and the final $y_{ij}$ to be obtained; $y_{ij}$ needs to answer $q_{ij}$ directly. |
| **The core method is actually used in the main analysis** | Specify the data that $K_j$ acts on in $d_i$ and how this step produces $y_{ij}$ or intermediate results required by the analysis. |
| **The inputs and relations required by the method have specific sources** | For each input in $\mathcal{U}_j$ and each relation in $\mathcal{E}_j$, specify the measurement, sample annotation or computational rule from which it comes, to avoid defining it during plan execution. |
| **The computational process from data to results is specified** | Specify how the data in $P$ are processed, which computations are performed in sequence and which choices affect $y_{ij}$. When multiple candidate parameters or construction approaches are involved, specify in advance how the final choice will be determined. |
| **The comparison can test the main result without circular evidence** | Specify which data or observations serve as $v_{ij}$ to test $y_{ij}$ and how they are compared. Final held-out observations cannot enter model fitting or selection. Null, sensitivity and method comparisons must specify the property they test and avoid treating a constructed target as independent evidence. |

A *P* that meets all five requirements is called a complete analysis plan. See Supplementary Note S2.2 for the boundaries and examples of completeness checking. If none of the initial candidates in Π*_ij_* is complete and some content can be added within *q_ij_* and *m_j_*, BioMINTREL selects a *P* to complete the specific missing parts. For example, it can add a computational rule for an intermediate quantity, specify an already permitted parameter choice, or clarify the split between test samples and fitting samples. Completion retains the *q_ij_*, *y_ij_*, analysis unit, *K_j_* and *C_ij_* of *P*. The maximum number of completion attempts is denoted by *R*. All five requirements are checked again after each completion attempt, and the process stops when a complete plan is obtained, or *R* attempts have been made. See Supplementary Note S2.3 for candidate selection, the scope of completion and stopping conditions.

When multiple candidates *P* are complete, BioMINTREL ranks them by the data basis of each input in *U_j_*, the link between *K_j_* and *y_ij_*, the specificity of *C_ij_* in *P*, and how fully the analysis choices have been determined. The four scores are denoted by *g_D_*(*P*), *g_M_* (*P*), *g_C_*(*P*) and *g_A_*(*P*), respectively, and each ranges from 0 to 1. Each score is calculated from the specific evidence in *P* according to fixed rules. For *P*, define

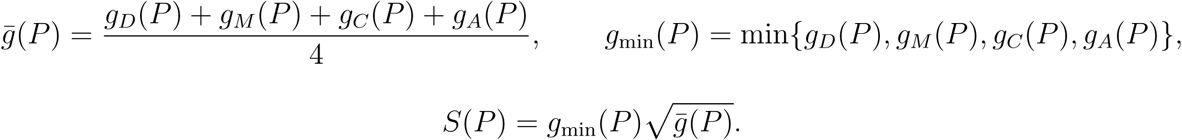

Here, *g̅*(*P*) is the mean of the four scores, *g*_min_(*P*) is the minimum score, and *S*(*P*) is the combined plan score. See Supplementary Note S2.4 and Supplementary Table S1 for score calculations and tie handling. For each pair (*t_i_, m_j_*), BioMINTREL retains the complete plan with the highest *S*(*P*), denoted by 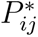, for the subsequent steps. If no candidate meets all five requirements within the completion budget, 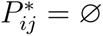.

### 5 Searching related studies and executing analyses

#### 5.1 SearchPriorArt: Searching studies related to the plan

BioMINTREL searches PubMed[56] for studies related to the final selected plan 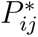. The search uses the data *d_i_*, method *m_j_*, question *q_ij_*, data representation *X_ij_* and main analysis content together. The relevant literature found is denoted by *L_ij_*. For studies in *L_ij_*, BioMINTREL compares whether they address the same question as *q_ij_*, use the same or a similar data representation to *X_ij_*, use the same or a similar method to *m_j_*, and whether 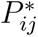 proposes a new analysis.

Based on this relation, the relation to the literature is denoted by *n_ij_* ∈ {N0, N1, N2, INSUFFICIENT}. N0 means that a direct precedent essentially the same as 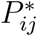 already exists; N1 means that existing studies use similar methods or analytical ideas, but the research question, data representation or specific analysis differs; N2 means that no close precedent was found within the scope of the search; INSUFFICIENT means that the current search results are insufficient to determine the relation between 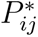 and existing studies. When 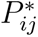 does not proceed to execution. When *n_ij_* is N1, N2 or INSUFFICIENT, 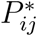 proceeds to the actual analysis. The relevant literature *L_ij_* and search judgment *n_ij_* are retained with 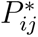.

### 5.2 ExecuteAndCheck: Performing computations and controls

At the execution stage, BioMINTREL uses the actual measurements and available sample annotations in *d_i_* to carry out the selected plan 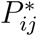. The plan defines the scientific *goal*, the main computation *K_j_* and the comparisons needed to assess the results. The main numerical results actually obtained are denoted by *Y_ij_*. The corresponding comparison results and uncertainty estimates are denoted by *V_ij_*.

To carry out each plan, BioMINTREL uses Codex as a coding agent. The agent receives a *goal* derived from the selected plan, together with the original dataset and method papers and access to the input data. The agent can also use dataset and database descriptions and method documentation, including repository README files, to clarify variable definitions, experimental conditions and method settings. The *goal* and plan vary across cases, but the same set of analysis *skills* is used throughout. These *skills* guide the agent in checking data and method requirements, preserving the core method, performing controls and assessing what the results support. The agent writes and revises the analysis code. After each revision, the code is run and the results are checked. Errors, missing results and incomplete comparisons guide the next revision. These revisions address problems in implementation without changing the scientific question or the core method. Researchers intervene when the available information is not enough to decide the next step. For the result visualization, the agent also generated the initial figures, and the authors subsequently adjusted their layout and visual appearance for presentation.

During analysis, BioMINTREL distinguishes three types of variables. The first consists of quantities directly measured in *d_i_*, such as gene expression levels, spatial coordinates or experimental conditions. The second consists of quantities obtained from measurements according to predefined rules, such as distances, adjacency relations or normalized expression values. The third consists of results calculated by *m_j_*, such as latent patterns, predictions or optimized allocations. Distinguishing these quantities clarifies whether each main result comes from original measurements, analysis definitions or method computations. See Supplementary Note S5, Supplementary Note S6, Supplementary Note S7 and Supplementary Note S8 for the case-specific constructions.

Comparisons follow the design specified in 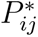 and address *q_ij_*. For predictive analyses, data used for fitting, selection and final validation are kept separate. Comparisons with existing methods use the same eligible data range and align the quantities at the level the methods share. A comparison can test the same output directly or examine whether a new representation adds information to an established summary. Permutation analysis shuffles only the relation being tested, while keeping other conditions such as sample numbers and group structure unchanged. The same computation is then repeated to test whether the observed result depends on that relation. Equal-count sampling compares samples or lineages with the same number of observations. This reduces the influence of differences in sampling counts. Parameter sensitivity analysis repeats the main computation over parameter ranges listed in advance to check whether the conclusion depends on a particular setting. See Supplementary Note S5, Supplementary Note S6, Supplementary Note S7 and Supplementary Note S8 for the case-specific computations and comparisons.

### 5.3 Interpret: Forming biological interpretations

BioMINTREL develops a biological interpretation *I_ij_* based on the actual results *Y_ij_* and comparison results *V_ij_*. The interpretation first states which new data relations or biological phenomena were observed, then which controls support this interpretation and which definitions and modeling assumptions of the data representation *X_ij_* the conclusion depends on. The relevant literature *L_ij_* is used to establish the link between *Y_ij_* and existing biological knowledge.

The scope of the interpretation is determined by the evidence supported by the actual computation. For example, if the results of a spatial model depend on how expression levels are converted into supply quantities and receiving limits, the conclusion needs to state this premise. If a perturbation analysis depends on constructed state relations, the results support the state connections observed under that representation and cannot be directly extended to untested causal relations. See case study 1, case study 2, case study 3 and case study 4 for the resulting biological interpretations.

For relations that still require further validation, BioMINTREL proposes corresponding follow-up measurements or experiments. The recommendations need to specify what to measure, which conditions to compare and which interpretations the experiment can distinguish. For example, independent tissue measurements can test predicted spatial relations, time-resolved data can test changes between adjacent states, or more balanced lineage sampling can assess whether the observed state is driven by a small number of cells.

## Evaluation

BioMINTREL aims to discover new data representations *X_ij_* and biological questions *q_ij_* (Method 3.1) from existing biological data *d_i_*, and develop these directions into executable analyses. This goal itself is open-ended: the same data can support several reasonable questions and representations, and a method from another field can also have several applications with supporting evidence, so it cannot be evaluated against a single standard answer. We examine whether the workflow is reliable from several complementary perspectives.

We evaluate BioMINTREL at the pipeline and case study levels. Pipeline evaluation uses two complementary sources of evidence. The first comes from expert consensus and assesses whether BioMINTREL’s judgments of task–method correspondence *C_ij_* (Method 3.1) agree with those of experienced human researchers. The second comes from related studies published in 2021 and later and examines whether pairs that the model considers suitable for further development more often correspond to method applications that appeared in later research. We then evaluate the role and performance of each module in BioMINTREL in turn. Finally, in four case studies, experts separately evaluate the analysis plans proposed by BioMINTREL and the results of the actual analyses.

### 1 Pipeline evaluation

We ran BioMINTREL separately with GPT-5.4 mini and Qwen3.6-35B as the underlying language model. For each model, we also evaluated a **direct LLM baseline**. The direct LLM receives the same task–method information as BioMINTREL, but assigns the Tier directly, without constructing the Method–data correspondence *C_ij_* or applying the correspondence-based Tier assignment rules (Method 3.1; Method 3.2). We refer to this direct judgment condition as the **direct LLM** below. We therefore compare direct GPT-5.4 mini with BioMINTREL (GPT-5.4 mini), and direct Qwen3.6-35B with BioMINTREL (Qwen3.6-35B).

#### 1.1 Evaluation of method-data correspondence using BioMDC-Bench

##### 1.1.1 Benchmark construction

BioMINTREL’s general workflow does not require users to specify a biological question or method name in advance, and the agent can identify potential questions and methods from the data paper and method paper. However, evaluating agreement between models and human experts requires comparing their judgments of the same pair. Here, we constructed BioMDC-Bench (Benchmark for Method–Data Correspondence in Biology) to evaluate whether a workflow can determine if the available biological data support the application of a candidate computational method to a specified biological task.

Specifically, we choose a use of the pipeline that is most convenient for pair-by-pair evaluation: specify the starting biological question *q_i_* and candidate method *m_j_* (Method 2.2) in advance, first form a clearly defined task–method pair (*t_i_, m_j_*) (Method 3.1), and then ask the model and experts to judge separately how the method can be applied in this pair. Two senior PhD students with 5 years of relevant research experience and one research scientist with 10 years of relevant research experience each independently assign a Tier according to the definitions in Method 3.2, using literature available in or before 2020. The final human reference retains only pairs for which all three experts give the same Tier. The resulting consensus benchmark contains 14 data papers, 19 tasks *t_i_*, and 12 candidate methods *m_j_*, giving 228 task–method pairs (see Evaluation 1.2).

The human reference and model predictions are independent of each other. The human experts do not view model predictions when making their judgments. The human Tier results are not inputs to BioMINTREL and do not participate in subsequent correspondence *C_ij_*, candidate ranking, or plan generation. Following the classification evaluation defined in Method 3.2, subsequent agreement evaluations consistently compare T1+T2 with T3. See Supplementary Note S3.1 for the final tasks, methods, and human reference labels.

##### 1.1.2 Comparison with direct LLMs

We directly compare BioMINTREL with the two underlying language models that it uses. The GPT-5.4 mini baseline and Qwen3.6-35B baseline each receive the same task-specific and method-specific information extracted from the dataset and method cards as BioMINTREL, but do not run BioMINTREL’s correspondence workflow *C_ij_* (Method 3.1), then directly judge the Tier. See Supplementary Note S3.2 for the model call settings.

We separately calculate the F1 score for agreement between each of the four groups of judgments and the human expert reference. For the same underlying model, we use a two-sided exact McNemar test[57, 58] to further compare the prediction results for each pair. See Supplementary Note S3.2 for the complete binary counts and test implementation. In addition to comparison with the expert reference, we directly compare whether the judgments of the two BioMINTREL implementations agree, recording intermodel agreement and Cohen’s *κ*[59]. See Supplementary Note S3.2 for the cross-tabulated counts.

#### 1.2 Evaluation using literature after 2020

##### 1.2.1 Evaluation evidence

To make pipeline evaluation less dependent on human Tiers, we additionally use studies published in 2021 and later as a second source of evidence. The later-literature evaluation still uses the task–method pair (*t_i_, m_j_*) (Method 3.1) as its unit and covers the same 228 pairs. For each pair, we collect information on whether method applications related to that combination appeared later. Related studies can use the same method, a similar method, or an analysis approach corresponding to that pair for the target task or a neighboring biological question, and later papers are not required to exactly match BioMINTREL’s complete correspondence *C_ij_* or final analysis plan 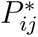 (Method 4.2). Each pair ultimately receives a binary later-literature outcome: a related study exists, or no related study was found in this collection. See Supplementary Note S3.3 for literature records for each pair, inclusion categories, and examples.

##### 1.2.2 Comparison between direct LLMs and BioMINTREL

For GPT-5.4 mini and Qwen3.6-35B, we separately read the Tiers already produced by the direct LLM and BioMINTREL. Each judgment approach divides the 228 pairs into T1+T2 and T3 according to its own results, and then matches them to the same later-literature outcome.

The direct LLM and BioMINTREL can place different numbers of pairs in T1+T2. We retain this difference because *how many combinations they are willing to take forward* is itself part of the two judgment approaches. We report the size of the T1+T2 group, the proportion of that group with later-literature evidence, and how many of all pairs with later-literature evidence are assigned to T1+T2.

We also compare the proportions with later-literature evidence in T1+T2 and T3 within each judgment approach, and report the risk ratio and 95% confidence interval. For each underlying model and each judgment approach, we separately use a two-sided Fisher’s exact test[60] to evaluate the association between Tier grouping and the later-literature outcome. To examine whether this relationship exists across different biological tasks, we calculate separately for each of the 19 tasks: the proportion with later-literature evidence in the model’s T1+T2 group for that task, and the proportion with later-literature evidence in its T3 group. See Supplementary Note S3.3 for the specific counts and statistical implementation.

We additionally compare the fixed expert reference with each BioMINTREL implementation on the 170 pairs with later-literature support. Coverage is the fraction assigned to T1+T2 within this subset. Agreement is the fraction for which the expert and model assignments coincide after combining T1 and T2. Two-sided exact McNemar tests compare the human-only and model-only T1+T2 counts. Holm correction treats the two human–model comparisons as one family[61]. We retain the original Tier of every pair and calculate their statistical significance (Supplementary Note S3.4).

### 2 Analysis of pipeline design

#### 2.1 Retrieval representation analysis

BioMINTREL’s regular retrieval uses both Scientific context and Analytical structure, and ranks candidate methods by the weighted score *r_ij_* in Method 2.2. To see clearly what information each of the two retrieval descriptions provides, we use two extreme settings of the same score formula in the evaluation: only Scientific context, with (*ω_c_, ω_a_*) = (1, 0); and only Analytical structure, with (*ω_c_, ω_a_*) = (0, 1). Analytical structure mainly helps find methods whose original application backgrounds differ but whose computational structure can correspond to the biological data, which are the hard-to-find pairs that BioMINTREL focuses on. From pairs with T2 in the consensus reference, we select the lowest quarter of Scientific context similarity 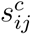 as the evaluation subset requiring retrieval across backgrounds. This stress-test subset meets two conditions at the same time: experts consider that the method can be used through constrained cross-disciplinary adaptation, but its original application background is not close to the target data. For each pair in the subset, we examine where its method ranks among the 12 candidate methods for the same task *t_i_*, record whether it enters the top three, and then compare the rank distributions and top-three coverage under the Scientific context and Analytical structure rankings. See Supplementary Note S3.5 for subset construction and ranking details.

#### 2.2 Scientific criteria ablation

We perform an ablation of the three scientific criteria in Method 3.3. For the Tier *τ_ij_* (Method 3.2) already obtained by BioMINTREL, we remove the three scientific criteria and use only the Tier to determine the priority of candidate pairs (*t_i_, m_j_*) for subsequent plan generation. Because each task *t_i_* ultimately allows only three methods to enter plan generation, we use the top-three reference support proportion as the main metric, that is, how many of these three limited places belong to T1+T2 in the consensus expert reference. This metric reflects the role of PrioritizeMethods: when a task has several applicable methods, which methods should be sent to subsequent plan generation first.

The evaluation ranking follows the basic logic of Method 3.3: T1 takes priority over T2, and T3 does not enter actual plan generation. In the condition with the scientific criteria, candidates with fewer clearly identified issues rank higher within the same Tier *τ_ij_*. See Supplementary Note S3.6 for the specific ordinal scores and the cap on the issue count used in the evaluation.

After the three scientific criteria are removed, several candidates often tie within the same Tier *τ_ij_*. To prevent the evaluation result from depending on an arbitrary method ID or a fixed tie-breaking order, we calculate the expected support count when ties occur at the top-three boundary. For example, if one place remains and two of four tied methods belong to human-reference T1+T2, that place contributes 0.5 expected supported combinations. As a reference without candidate ranking information, we also calculate the expected reference support proportion when three methods are selected uniformly without replacement from the 12 methods for each task. See Supplementary Note S3.6 for the complete ranking scores.

#### 2.3 Repeated Formulation analysis

Each selected task–method pair (*t_i_, m_j_*)*, m_j_* ∈ S*_i_* (Method 3.3) independently generates the 5 candidate plans in Π*_ij_* (Method 4.1), and we use these 5 plans to evaluate the effect of repeated generation. The single-Formulation condition checks only whether the first candidate 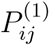 is complete, whereas the five-Formulations condition checks whether at least 1 of the 5 candidates in Π*_ij_* is complete. For example, when the first candidate 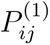 does not pass the completeness check but the third candidate 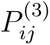 does, that pair is recorded as incomplete in the single-generation condition and complete in the five-generation condition. We thus compare whether increasing the number of independent generations increases the chance of obtaining a complete analysis plan for the same task–method pair (*t_i_, m_j_*).

Each of the five candidates in Π*_ij_* uses the same inputs without access to the other candidates. The evaluation does not combine content from several incomplete candidates into one plan, nor select one of the five after generation as the “single-generation” result. All candidates *P* are assessed using the five completeness requirements in Method 4.2. We summarize results at the pair (*t_i_, m_j_*) and task *t_i_* (Method 3.1) levels. Pair completeness is the proportion of selected pairs that produce at least one complete plan *P*, and the same pair is counted only once even if it has several complete candidates. Task coverage is the proportion of all tasks for which at least one selected method produces a complete plan. A task with several complete plans is also counted only once. The former assesses whether a specific task–method correspondence *C_ij_* can be developed into a plan, and the latter assesses whether a starting biological question *q_i_* already has at least one executable analysis.

The three pairs for the same task *t_i_* share data *d_i_* and a starting question *q_i_* (Method 3.1), so uncertainty in pair completeness is estimated by resampling tasks[62]. In each iteration, tasks are sampled with replacement, and the three pairs corresponding to each task are retained together, with the 2.5th and 97.5th percentiles of the resampling distribution forming the 95% confidence interval after 5,000 repetitions. See Supplementary Note S3.7 for generation records, completeness judgments, and resampling settings.

### 3 Case-study evaluation

One senior PhD student with 5 years of relevant research experience and one research scientist with 10 years of relevant research experience participate in the case study evaluation. The four cases are evaluated in two stages: the two human experts first compare analysis plans without knowing their sources, and then evaluate the actual analyses completed by BioMINTREL. The first stage evaluates whether the research direction itself is worth pursuing, and the second evaluates whether the actual results, validation, and interpretations are reasonable.

#### 3.1 Blinded evaluation of analysis plans

##### 3.1.1 Analysis approaches preparation

Three analysis plans addressing the same biological question are prepared for each case. The first is the plan 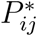 (Method 4.2) generated and ultimately selected by BioMINTREL. The second uses established analysis methods commonly used in the target field, representing further analysis of the same data within that field’s existing methods and serving to compare whether BioMINTREL and established specialized methods can propose different but reasonable analysis directions. The third does not introduce a new cross-disciplinary method or new data representation, and directly uses existing measurements, groups, and annotations for conventional statistical comparisons, such as comparisons of group differences, data distributions, correlations, or relationships between existing annotations, with the aim of providing a more basic reference to judge whether the new method transfer and data representation actually add analysis content.

The three plans use the same data *d_i_* (Method 1) and a common review question, but each retains its own analysis path. Each plan provides experts with the question to be answered, the main analysis approach, expected results, and planned validation. Reviewers judge this scientific content without using BioMINTREL’s internal Tier *τ_ij_* (Method 3.2), correspondence *C_ij_* (Method 3.1), or completeness score. See Supplementary Note S4.1 for the case correspondence and assessment dimensions.

##### 3.1.2 Plan review settings

Before the review begins, the two human experts are unfamiliar with BioMINTREL’s workflow and do not know which of the three plans was generated by BioMINTREL. The plans for each case are labeled A, B, and C and presented in a shuffled order. The actual analysis results are also withheld from the experts, so this stage evaluates only the plans themselves and is not influenced by how good or poor the final results are.

Experts first evaluate each of the three plans for analysis novelty, biological relevance, methodological reliability, the match between the method and the question, and whether the validation design is sufficient to test the main interpretation. Each dimension uses a 1–4 scale, with higher scores indicating a more positive evaluation; see Supplementary Note S4.1 and Supplementary Data 3 (`plan-review forms`) for the assessment dimensions and individual scores.

After scoring the individual items, experts give an overall ranking of the three plans and select the plan they consider most worth executing. The review form also separately asks whether each plan is worth pursuing and which plan is most likely to provide additional information. Experts also record the validation or analysis they would most like to add and their confidence in their own judgment.

One expert’s complete review of one case is recorded as one reviewer–case assessment. Scores for each type of plan are summarized separately across the five dimensions, together with the overall first choice, recommendations to proceed, and the choice “most likely to provide additional information.” See Supplementary Note S4.2 for the complete score tables, rankings, and open comments.

#### 3.2 Evaluation of completed analyses

After the blinded plan review, the two experts further read the actual analyses completed by BioMINTREL in the four cases and assess whether the results, validation, and interpretations ultimately obtained by BioMINTREL are reasonable. Experts receive an analysis summary, main results *Y_ij_*, completed controls or validation *V_ij_* (Method 5.2), and the corresponding biological interpretation for each case.

Experts separately evaluate five aspects: whether the results are relevant to the target biological question; whether the data representation *X_ij_* (Method 3.1) and analysis framework actually used are suitable for answering that question; whether the methods and statistical treatment are reasonable; whether the available evidence and validation are sufficient to support the main interpretation; and whether the analysis adds new information or a further level of explanation relative to conventional analysis. Each dimension uses a 1–4 scale.

Experts also judge whether the analysis is worth further study, identify its most valuable part, and state one analysis or measurement they would most like to add. Each assessment also records the expert’s confidence in their own judgment. Completed analyses are still summarized by reviewer–case assessment, giving eight assessments in total. The five scores are summarized separately, together with recommendations for further research and open comments. See Supplementary Note S4.3 for the individual scores, confidence ratings and comments.

## Data availability

Case study 1 uses deep mutational scanning data from the study of antifolate resistance in Pneumocystis jirovecii dihydrofolate reductase (https://doi.org/10.1371/journal.pgen.1011252). Raw sequencing reads are available under BioProject accession PRJNA1013557 (https://www.ncbi.nlm.nih.gov/bioproject/PRJNA1013557). Processed variant measurements are available at https://github.com/Landrylab/Rouleau_et_al_2024.

Case study 2 uses the 10x Genomics Fresh Frozen Visium on CytAssist Human Breast Cancer dataset, available at https://www.10xgenomics.com/datasets/fresh-frozen-visium-on-cytassist-human-breast-cancer-probe-based-whole-transcriptome-profiling-2-standard.

Case study 3 uses unfolded protein response Perturb-seq data (https://doi.org/10.1016/j.cell.2016.11.048), available from the Gene Expression Omnibus under accession GSE90546 (https://www.ncbi.nlm.nih.gov/geo/query/acc.cgi?acc=GSE90546), including sample GSM2406681 (https://www.ncbi.nlm.nih.gov/geo/query/acc.cgi?acc=GSM2406681).

Case study 4 uses data from the Watermelon lineage-tracing study (https://doi.org/10.1038/s41586-021-03796-6), available from the Gene Expression Omnibus under accession GSE150949 (https://www.ncbi.nlm.nih.gov/geo/query/acc.cgi?acc=GSE150949). The original study data and analysis code are available at https://github.com/yaaraore/Watermelon.

Supplementary Data 1 contains the 228 BioMDC-Bench task–method pairs, their source publications, expert consensus Tiers, later-literature evidence and saved model decisions. Supplementary Data 2 contains the saved pipeline outputs and available case-study plans. Supplementary Data 3 contains the original expert review documents for the four case studies.

## Code availability

Code to reproduce the pipeline, benchmark and analytical results is accessible at: https://github.com/7DongJiaqi/BioMINTREL.

## References

1. Cao, J. et al. The single-cell transcriptional landscape of mammalian organogenesis. Nature 566, 496–502 (2019).

2. Heumos, L., Schaar, A. C., Lance, C. et al. Best practices for single-cell analysis across modalities. Nature Reviews Genetics 24, 550–572 (2023).

3. Trapnell, C. Revealing gene function with statistical inference at single-cell resolution. Nature Reviews Genetics 25, 623–638 (2024).

4. Lu, C. et al. The AI scientist: Towards fully automated open-ended scientific discovery. arXiv preprint arXiv:2408.06292 (2024). URL https://arxiv.org/abs/2408.06292.

5. Schmidgall, S. et al. Agent laboratory: Using LLM agents as research assistants. In Findings of the Association for Computational Linguistics: EMNLP 2025, 5977–6043 (Association for Computational Linguistics, 2025). URL https://aclanthology.org/2025.findings-emnlp.320/.

6. Yamada, Y. et al. The AI scientist-v2: Workshop-level automated scientific discovery via agentic tree search. arXiv preprint arXiv:2504.08066 (2025). URL https://arxiv.org/abs/2504.08066.

7. Schiebinger, G. et al. Optimal-transport analysis of single-cell gene expression identifies developmental trajectories in reprogramming. Cell 176, 928–943.e22 (2019).

8. Burge, C. & Karlin, S. Prediction of complete gene structures in human genomic DNA. Journal of Molecular Biology 268, 78–94 (1997).

9. Pevzner, P. A., Tang, H. & Waterman, M. S. An eulerian path approach to DNA fragment assembly. Proceedings of the National Academy of Sciences of the United States of America 98, 9748–9753 (2001).

10. Zerbino, D. R. & Birney, E. Velvet: Algorithms for de novo short read assembly using de Bruijn graphs. Genome Research 18, 821–829 (2008).

11. Ghareeb, A. E. et al. A multi-agent system for automating scientific discovery. Nature 655, 497–505 (2026).

12. Swanson, K., Wu, W., Bulaong, N. L., Pak, J. E. & Zou, J. The virtual lab of AI agents designs new SARS-CoV-2 nanobodies. Nature 646, 716–723 (2025).

13. Alber, S. et al. CellVoyager: AI CompBio agent generates new insights by autonomously analyzing biological data. Nature Methods 23, 749–759 (2026).

14. Huang, K., et al. Biomni: A general-purpose biomedical AI agent. bioRxiv 2025.05.30.656746 (2025).

15. Cao, L., et al. BiOmics: A foundational agent for grounded and autonomous multi-omics interpretation. bioRxiv 2026.01.17.699830 (2026).

16. Shen, A., Druckmann, S. & Zou, J. Unlocking LLM creativity in science through analogical reasoning. arXiv preprint arXiv:2605.11258 (2026).

17. Knapp, B. et al. Ten simple rules for a successful cross-disciplinary collaboration. PLOS Computational Biology 11, e1004214 (2015). URL 10.1371/journal.pcbi.1004214.

18. Gentner, D. Structure-mapping: A theoretical framework for analogy. Cognitive Science 7, 155–170 (1983). URL 10.1207/s15516709cog0702_3.

19. OpenAI. GPT-5.4 Mini Model (2026). URL https://developers.openai.com/api/docs/models/gpt-5.4-mini. Accessed 7 October 2026.

20. Qwen Team. Qwen3.6-35B-A3B: Model card (2026). URL https://huggingface.co/Qwen/Qwen3.6-35B-A3B. Accessed 7 October 2026.

21. Rouleau, F. D. et al. Deep mutational scanning of Pneumocystis jirovecii dihydrofolate reductase reveals allosteric mechanism of resistance to an antifolate. PLOS Genetics 20, e1011252 (2024).

22. Rodrigues, J. V. et al. Biophysical principles predict fitness landscapes of drug resistance. Proceedings of the National Academy of Sciences of the United States of America 113, E1470–E1478 (2016).

23. Tareen, A. et al. MAVE-NN: learning genotype-phenotype maps from multiplex assays of variant effect. Genome Biology 23, 98 (2022).

24. 10x Genomics. Fresh frozen Visium on CytAssist: Human breast cancer, probe-based whole transcriptome profiling. 10x Genomics dataset (2023). URL https://www.10xgenomics.com/datasets/fresh-frozen-visium-on-cytassist-human-breast-cancer-probe-based-whole-transcriptome-profiling-2-standard. Accessed 18 September 2026.

25. Lücken, L., Mitschke, N., Dittmar, T. & Blasius, B. Network flow methods for NMR-based compound identification. Analytical Chemistry 97, 4832–4840 (2025).

26. Ahuja, R. K., Magnanti, T. L. & Orlin, J. B. Network Flows: Theory, Algorithms, and Applications (Prentice Hall, 1993).

27. Cang, Z. et al. Screening cell-cell communication in spatial transcriptomics via collective optimal transport. Nature Methods 20, 218–228 (2023).

28. Jin, S. et al. Inference and analysis of cell-cell communication using CellChat. Nature Communications 12, 1088 (2021).

29. Müller, A., et al. Involvement of chemokine receptors in breast cancer metastasis. Nature 410, 50–56 (2001).

30. Liu, F., Zhang, J., Gu, X., Guo, Q. & Guo, W. Single-cell transcriptome sequencing reveals SPP1-CD44-mediated macrophage-tumor cell interactions drive chemoresistance in TNBC. Journal of Cellular and Molecular Medicine 28, e18525 (2024).

31. Huang, L., Qin, S., Xia, L., Ma, F. & Chen, L. Integrative single-cell sequencing analysis distinguishes survival-associated cells from the breast cancer microenvironment. Cancer Medicine 12, 12896–12911 (2023).

32. Pellegrino, B. et al. CD74 promotes the formation of an immunosuppressive tumor microenvironment in triple-negative breast cancer in mice by inducing the expansion of tolerogenic dendritic cells and regulatory B cells. PLOS Biology 22, e3002905 (2024).

33. Li, H. et al. Decoding functional cell-cell communication events by multi-view graph learning on spatial transcriptomics. Briefings in Bioinformatics 24, bbad359 (2023).

34. Wang, X. et al. Spatially resolved atlas of breast cancer uncovers intercellular machinery of venular niche governing lymphocyte extravasation. Nature Communications 16, 3348 (2025).

35. Rydén, L., et al. Tumor specific VEGF-A and VEGFR2/KDR protein are co-expressed in breast cancer. Breast Cancer Research and Treatment 82, 147–154 (2003).

36. Zhang, W., Ran, S., Sambade, M., Huang, X. & Thorpe, P. E. A monoclonal antibody that blocks VEGF binding to VEGFR2 (KDR/Flk-1) inhibits vascular expression of Flk-1 and tumor growth in an orthotopic human breast cancer model. Angiogenesis 5, 35–44 (2002).

37. Adamson, B. et al. A multiplexed single-cell CRISPR screening platform enables systematic dissection of the unfolded protein response. Cell 167, 1867–1882.e21 (2016).

38. Metzner, P., Schütte, C. & Vanden-Eijnden, E. Transition path theory for markov jump processes. Multiscale Modeling & Simulation 7, 1192–1219 (2009).

39. E, W. & Vanden-Eijnden, E. Towards a theory of transition paths. Journal of Statistical Physics 123, 503–523 (2006).

40. E, W. & Vanden-Eijnden, E. Transition-path theory and path-finding algorithms for the study of rare events. Annual Review of Physical Chemistry 61, 391–420 (2010).

41. Iida, Y. et al. SEL1L protein critically determines the stability of the HRD1-SEL1L endoplasmic reticulum-associated degradation (ERAD) complex to optimize the degradation kinetics of ERAD substrates. Journal of Biological Chemistry 286, 16929–16939 (2011).

42. Bertolotti, A., Zhang, Y., Hendershot, L. M., Harding, H. P. & Ron, D. Dynamic interaction of BiP and ER stress transducers in the unfolded-protein response. Nature Cell Biology 2, 326–332 (2000).

43. Sundaram, A., Plumb, R., Appathurai, S. & Mariappan, M. The Sec61 translocon limits IRE1alpha signaling during the unfolded protein response. eLife 6, e27187 (2017).

44. Wang, L. et al. UFMylation of RPL26 links translocation-associated quality control to endoplasmic reticulum protein homeostasis. Cell Research 30, 5–20 (2020).

45. Oren, Y. et al. Cycling cancer persister cells arise from lineages with distinct programs. Nature 596, 576–582 (2021).

46. Rockafellar, R. T. & Uryasev, S. Optimization of conditional value-at-risk. The Journal of Risk 2, 21–41 (2000).

47. Rockafellar, R. T. & Uryasev, S. Conditional value-at-risk for general loss distributions. Journal of Banking & Finance 26, 1443–1471 (2002).

48. An, Y., Liang, J., Schild, S. E., Bues, M. & Liu, W. Robust treatment planning with conditional value at risk chance constraints in intensity-modulated proton therapy. Medical Physics 44, 28–36 (2017).

49. Shaffer, S. M. et al. Rare cell variability and drug-induced reprogramming as a mode of cancer drug resistance. Nature 546, 431–435 (2017).

50. Emert, B. L. et al. Variability within rare cell states enables multiple paths toward drug resistance. Nature Biotechnology 39, 865–876 (2021).

51. McInnes, L., Healy, J. & Melville, J. UMAP: Uniform manifold approximation and projection for dimension reduction. arXiv preprint arXiv:1802.03426 (2018). URL https://arxiv.org/abs/1802.03426.

52. National Center for Biotechnology Information. PMC Open Access Subset (2026). URL https://pmc.ncbi.nlm.nih.gov/tools/openftlist/. Accessed 18 September 2026.

53. Lee, D. D. & Seung, H. S. Learning the parts of objects by non-negative matrix factorization. Nature 401, 788–791 (1999).

54. Singh, A., D’Arcy, M., Cohan, A., Downey, D. & Feldman, S. SciRepEval: A multi-format benchmark for scientific document representations. Proceedings of the 2023 Conference on Empirical Methods in Natural Language Processing 5548–5566 (2023). Association for Computational Linguistics, Singapore.

55. Cawley, G. C. & Talbot, N. L. C. On over-fitting in model selection and subsequent selection bias in performance evaluation. Journal of Machine Learning Research 11, 2079–2107 (2010). URL https://jmlr.org/papers/v11/cawley10a.html.

56. National Center for Biotechnology Information. About PubMed (2026). URL https://pubmed.ncbi.nlm.nih.gov/about/. Accessed 7 October 2026.

57. McNemar, Q. Note on the sampling error of the difference between correlated proportions or percentages. Psychometrika 12, 153–157 (1947).

58. Fagerland, M. W., Lydersen, S. & Laake, P. The McNemar test for binary matched-pairs data: mid-p and asymptotic are better than exact conditional. BMC Medical Research Methodology 13, 91 (2013).

59. Cohen, J. A coefficient of agreement for nominal scales. Educational and Psychological Measurement 20, 37–46 (1960).

60. Fisher, R. A. The logic of inductive inference. Journal of the Royal Statistical Society 98, 39–54 (1935).

61. Holm, S. A simple sequentially rejective multiple test procedure. Scandinavian Journal of Statistics 6, 65–70 (1979). URL https://www.jstor.org/stable/4615733.

62. Field, C. A. & Welsh, A. H. Bootstrapping clustered data. Journal of the Royal Statistical Society: Series B (Statistical Methodology) 69, 369–390 (2007).

63. Rao, S. S. P. et al. A 3D map of the human genome at kilobase resolution reveals principles of chromatin looping. Cell 159, 1665–1680 (2014).

64. Cuturi, M. Sinkhorn distances: Lightspeed computation of optimal transport. In Advances in Neural Information Processing Systems 26, 2292–2300 (2013).

65. Zeira, R., Land, M., Strzalkowski, A. & Raphael, B. J. Alignment and integration of spatial transcriptomics data. Nature Methods 19, 567–575 (2022).

66. Liu, X., Zeira, R. & Raphael, B. J. Partial alignment of multislice spatially resolved transcriptomics data. Genome Research 33, 1124–1132 (2023).

67. Chang, S. et al. PertSpectra: Interpretable matrix factorization for predicting functional impact of genetic perturbation experiments. In Pacific Symposium on Biocomputing, vol. 31, 465–479 (World Scientific, 2026). URL https://psb.stanford.edu/psb-online/proceedings/psb26/chang_s.pdf.

68. Benjamini, Y. & Hochberg, Y. Controlling the false discovery rate: A practical and powerful approach to multiple testing. Journal of the Royal Statistical Society: Series B (Methodological) 57, 289–300 (1995).

69. Delgado, J., Radusky, L. G., Cianferoni, D. & Serrano, L. FoldX 5.0: working with RNA, small molecules and a new graphical interface. Bioinformatics 35, 4168–4169 (2019).

70. Kruskal, W. H. & Wallis, W. A. Use of ranks in one-criterion variance analysis. Journal of the American Statistical Association 47, 583–621 (1952).

71. Mann, H. B. & Whitney, D. R. On a test of whether one of two random variables is stochastically larger than the other. The Annals of Mathematical Statistics 18, 50–60 (1947).

72. Kingma, D. P. & Ba, J. Adam: A method for stochastic optimization. In International Conference on Learning Representations (2015). URL https://arxiv.org/abs/1412.6980.

73. CellChat developers. CellChat analysis of spatial transcriptomics data. Official software tutorial. URL https://github.com/jinworks/CellChat/blob/main/tutorial/CellChat_analysis_of_spatial_transcriptomics_data.Rmd. Accessed 7 October 2026.

74. Traag, V. A., Waltman, L. & van Eck, N. J. From Louvain to Leiden: guaranteeing well-connected communities. Scientific Reports 9, 5233 (2019).

75. Noé, F., Schütte, C., Vanden-Eijnden, E., Reich, L. & Weikl, T. R. Constructing the equilibrium ensemble of folding pathways from short off-equilibrium simulations. Proceedings of the National Academy of Sciences 106, 19011–19016 (2009).

76. Helfmann, L., Ribera Borrell, E., Schütte, C. & Koltai, P. Extending transition path theory: Periodically driven and finite-time dynamics. Journal of Nonlinear Science 30, 3321–3366 (2020).

77. Nichols, T. E. & Holmes, A. P. Nonparametric permutation tests for functional neuroimaging: A primer with examples. Human Brain Mapping 15, 1–25 (2002).

