## Supplementary Note for "BioMINTREL expands biological problem solving through cross-disciplinary representations"

**Accompanying data files.**

**Supplementary Data 1** (`Supplementary_Data_1.xlsx`): benchmark tasks, methods, reference labels, literature records and saved model decisions.

**Supplementary Data 2** (`Supplementary_Data_2.zip`): benchmark inputs, saved pipeline outputs and available case-study plans.

**Supplementary Data 3** (`Supplementary_Data_3.zip`): original expert reviews of candidate plans and completed analyses for four case studies.

**Contents.**


S1, Boundaries in method–data correspondence

S2, Completing and selecting analysis plans

S3, Details of the benchmark comparisons

S4, Expert review materials and interpretation of scores

S5, Computations for the PjDHFR dose-response analysis

S6, Definitions and comparisons in spatial allocation

S7, Constructing and comparing perturbation response routes

S8, Sampling and prediction in the lineage analysis

**Supplementary Note S1: Boundaries in method–data correspondence**

### S1.1 Distinguishing the four construction statuses

*Observed* and *deterministic* differ in whether the required input or relationship already exists in the data. A recorded coordinate or treatment label is *observed*. A distance calculated from coordinates or a response score calculated from expression is *deterministic* when a fixed rule uniquely determines it from the recorded measurements. If that rule has not yet been specified, the required quantity is *missing/open*.

For a spatial method that requires a connected, undirected graph, consider three candidates: 5, 10 or 20 nearest neighbours per location. Each graph uses Euclidean distance, connects two locations when either selects the other, assigns unit edge weights and resolves distance ties by a fixed location identifier. The selection rule chooses the smallest of the three neighbour counts that connects all locations. If the neighbour count is fixed at 10 before analysis, spatial adjacency is *deterministic*. If the analysis chooses among 5, 10 and 20 neighbours by the connectivity rule above, spatial adjacency is *bounded*. The selection algorithm itself is *deterministic*, but the construction is still *bounded* because the definition of adjacency must be chosen from three predefined candidates. If none of the three graphs is connected, none satisfies the method's stated requirement. Continuing the analysis would then require a new graph definition outside the predefined candidate set, so the required adjacency relation remains *missing/open*. A *bounded* construction needs both a finite candidate set and a prespecified rule for choosing among the candidates. Listing 5, 10 and 20 without a selection rule is therefore *missing/open*. "Choose a suitable neighbourhood" is also *missing/open* because neither the allowed range nor the selection rule is fixed. Choosing the graph with the strongest association with the final outcome is not an allowed *bounded* choice because the outcome being tested would also determine the graph used in that test.

For a flow model, simply assigning positive and negative values to nodes is not enough to define biological supply and demand. The required quantity is the amount that each node contributes to or receives from the flow, so the correspondence must define which nodes act as sources or sinks, what the sign means biologically, and how the magnitude and units follow from the measurements. If the data directly record the amount supplied or demanded by each node, this requirement is *observed*. If the data record a related quantity, such as expression, and a fixed rule converts that measurement into node supply or demand with the source–sink roles, sign convention, scale and units specified in advance, the requirement is *deterministic*. If several explicit mappings from the measurements to supply or demand are allowed and a prespecified criterion selects one without using the final biological result, the requirement is *bounded*. If the analysis only assigns positive values to some nodes and negative values to others, or leaves the expression-to-flow mapping to be chosen later, without defining these biological roles and quantities, the requirement remains *missing/open*.

### S1.2 Preserving the core and obtaining the biological result

Core preservation asks whether the transferred analysis still performs the defining calculation of the source method under its defining constraints. For non-negative matrix factorization, changing the number of factors can preserve the core if the analysis still factorizes the matrix into non-negative factors under the original objective[53]. Replacing this factorization with principal-component analysis changes the defining calculation and therefore does not preserve the core. Merely referring to the step as non-negative matrix factorization is also insufficient. The plan must actually perform that factorization.

What must happen after factorization depends on the biological question. If the question asks how strongly shared expression programmes contribute to each sample, the programme weights are already the main result. If the question asks whether treatments differ, the weights alone do not answer it: the plan must define the treatment groups and the comparison between their weights. If the question asks for expression in a new condition, the plan must also explain how to obtain that condition's programme weights. Rescaling an output back to its original measurement units is only a unit conversion and does not create a new biological result. Section S2.2 gives the new-condition example.

Executing the source method and using its output are separate requirements. A plan can fit the factorization correctly and then discard the factors while another model produces the final result. In that case the method core is retained, but it contributes nothing to the proposed biological comparison. The core-contribution criterion checks whether the transferred method's output actually enters the main result.

### S1.3 Changes that distinguish T1, T2 and T3

Each required input and relationship must be defined separately. The same measurement can support more than one requirement, but each use needs its own construction. For example, spatial coordinates can provide both inter-location distances and graph adjacency. A distance formula defines the distances. The adjacency relation still needs a rule that states which locations are connected. Specifying the distance twice leaves the adjacency relation undefined.

The same data can support different Tiers when the biological question changes. Consider a non-negative expression matrix and non-negative matrix factorization. If the starting question asks which shared expression programmes describe the samples, the matrix can support a direct T1 application when the other T1 conditions are satisfied. If the starting question concerns perturbation responses and the analysis reformulates it as how perturbations change the weights of shared programmes, the inputs can remain unchanged while the pair becomes T2 because the biological question has been reformulated within a defined scope. If the new question has no clear connection to the starting question, the correspondence is T3.

A unit conversion is a minor adjustment when it preserves the same analytical object. For example, converting spatial coordinates from micrometres to millimetres and converting the distance threshold by the same factor leaves the graph unchanged. If the coordinates are converted and the threshold keeps its original numerical value, the set of connected locations changes. The adjacency relation has changed and needs to be assessed as a new construction.

The Tier depends on what a bounded choice changes. A prespecified choice among component numbers within the source matrix-factorization method can remain T1 when it changes only an internal method parameter and leaves the required inputs, data relations, biological question and overall adaptation unchanged. In contrast, choosing a neighbourhood size from $\{5, 10, 20\}$ changes the adjacency relation supplied to a graph method. That required relationship is therefore bounded, and the pair is T2 when the remaining correspondence conditions are satisfied.

Tier assignment asks whether the method–data correspondence is sufficiently defined to support the proposed transfer. It does not depend on the eventual effect size, prediction accuracy or statistical significance. A pair can therefore receive T1 or T2 before any result has been observed. An undefined input or relationship required by the core calculation instead leaves the correspondence incomplete and gives T3.

Tier and plan completeness are separate decisions. T1 or T2 means that the required correspondence between the method and the data has been established. A complete plan additionally specifies how the method output produces the final biological result, which observations enter fitting and evaluation, and which comparison tests that result. Supplementary Note S2 describes these additional requirements.

#### S1.4 Applying the three scientific criteria

**Biological basis.** A required quantity can be mathematically well defined but still lack a biological basis for the role assigned to it. For example, expression distance between two cell states can be calculated directly from their expression vectors. If a transport model uses this distance as its cost, the analysis must also explain why larger expression differences should represent greater difficulty or lower plausibility of moving between those states. A formula alone establishes how the cost is calculated. It does not establish why that cost is biologically meaningful for the question being studied. This criterion checks whether each constructed input or relation has both a clear calculation and a biological justification for the role it plays in the method.

**Core contribution.** Running the transferred method is not enough. Its output must be used to produce the main result. For example, factorization contributes to a response prediction when the resulting factor weights enter the prediction model. If the analysis obtains the full prediction from another calculation and does not use the factorization output, then the factorization adds no computational contribution to that result. This criterion therefore asks a direct question: does the transferred method produce information that the main analysis actually uses? It does not ask whether another method could also solve the same task.

**Validation.** The proposed comparison must test the specific claim made from the main result using evidence that did not determine that result. For example, if cell-type labels are used to choose the embedding that best separates those cell types, separation measured with the same labels cannot provide an independent test because those labels already determined which embedding was selected[55]. Labels or observations kept outside that selection can test whether the selected embedding reproduces the same structure. For a null comparison, the shuffle should break the relationship that produced the claimed result. The comparison then asks whether the observed result exceeds what remains after that relationship is broken. For a sensitivity analysis, the analysis should vary the specific setting whose influence is in question and test whether the conclusion depends on that setting. This criterion therefore asks whether the proposed comparison can actually challenge the claim made from the result.

Each scientific criterion contributes either zero or one issue to candidate ranking. Multiple problems of the same type do not receive additional penalties. For example, several unsupported biological interpretations still give one biological-basis issue. When the available information is insufficient to decide whether a criterion is violated, BioMINTREL does not count a definite issue. Section S3.6 gives the scores used in the ranking comparison.

#### S1.5 Forming tasks and models from multiple sources

Input papers can contribute different parts of a task or model. Across datasets, BioMINTREL identifies which observations can be related, for example through shared samples, complementary measurements or linked biological stages, and uses those relations to define a joint task. Across methods, it considers complementary computations, sequential input–output relations, comparisons and suitable combinations of components or ideas. The resulting formulation states what each source contributes and how those contributions address the biological question.

The spatial case jointly considered mcfNMR, Alignstein and CINDA. The saved plan assigned mcfNMR the primary role in capacity-constrained minimum-cost flow and unmatched-supply handling, used Alignstein to inform local correspondence and geometric costs, and drew on CINDA for an auxiliary view of global association and factorized costs. These distinct roles shaped a common spatial allocation model (Supplementary Data 2 (`spatial case inputs and saved plan`)).

The perturbation case jointly considered PyEMMA 2, PCCA+ and transition-path theory. Their contributions concerned state-space organization, spectral coarse-graining and source-to-target path calculations, respectively. The saved plan related these operations through a shared response-state graph, and the analysis computed path statistics and derived a coarser state grouping on that representation. The 11 expression-defined states used in the main analysis were obtained by Leiden clustering. The coarser grouping summarized relations among those states. Together, the spatial and perturbation cases illustrate how several method papers can inform one analysis, with distinct contributions to its representation and computations (Supplementary Data 2 (`perturbation case inputs and saved plan`)).

#### S1.6 Using information from figures, supplementary materials and documentation

In the PjDHFR analysis, variant-effect tables were combined with FoldX structural scores supplied with the source study's figures[21]. In the perturbation analysis, gene sets drawn from Fig. 3F of the source study and its accompanying text were used to calculate UPR scores. Annotations of perturbation groups and their responses informed comparisons across the inferred routes[37].

In the lineage analysis, pathway findings in Watermelon's main and Extended Data figures guided the choice of candidate programmes and biological controls[45]. The supplied MAVE-NN materials also included the repository's README description of genotype–phenotype modelling, while the data description for the spatial case identified the Visium section and its measurements (Supplementary Data 2 (`case input records`)).

### Supplementary Note S2: Completing and selecting analysis plans

#### S2.1 Settings for initial generation and completion

Each selected task–method pair receives five independent initial plans ($J = 5$). These plans explore alternative formulations of the same established method–data correspondence and do not see one another. If none of the five plans is complete, BioMINTREL can make at most two targeted completion attempts ($R = 2$), but only on candidates whose missing content can be supplied without changing the biological question, main result, analysis unit, core method or established method–data correspondence. Both language models use temperature 0.3 for initial generation, 0.2 for completion and 0.0 for completeness assessment. The repeated-generation evaluation in Section S3.7 uses only the five independent initial plans, and targeted completion is excluded from that comparison.

#### S2.2 When an apparently runnable analysis is still incomplete

A plan is incomplete when an unresolved step remains between the available data, the transferred method and the biological result that the plan claims to test. Producing an intermediate representation is not enough if the plan does not specify how that representation produces the main result. A plan is also incomplete when a quantity that determines the analysis still has several scientifically different definitions and the plan has not selected one in advance. The examples below illustrate these boundaries.

**Predicting expression after matrix factorization.** Let $X_{\text{train}} \approx W_{\text{train}} H$, where rows of $X_{\text{train}}$ are training conditions, rows of $H$ are shared gene programmes, and each row of $W_{\text{train}}$ contains one training condition's programme weights. Estimating $H$ completes the factorization of the training data, but it does not define a prediction for a new condition. The new condition requires weights $w_*$ before the plan can calculate

$$\hat{x}_* = w_* H.$$

A complete prediction plan must therefore specify what information is available for that condition before its target expression is known and how that information produces $w_*$.

Using the new condition's measured target expression to fit $w_*$ does not solve this prediction problem. It reconstructs an expression profile that was already used to estimate the weights. A held-out prediction requires $w_*$ to be obtained without using the target expression that will later evaluate $\hat{x}_*$.

**A transport analysis of Hi-C contacts.** A plan is also incomplete when a quantity that determines the calculation still has several scientifically different definitions and the plan leaves that choice unresolved. Hi-C measurements count spatial contacts between genomic regions[63]. A transport calculation requires two input distributions, a cost between regions and a defined result from the transport solution[64]. Writing "genomic distance or a transformation" does not define the cost, because raw distance and different nonlinear transformations assign different relative penalties to short and long genomic separations. The plan must choose the formula, units and scale before execution and use the same definition throughout the analysis.

For held-out contact prediction, the test target must remain outside every step that can influence the prediction, not only the final model fit. The plan must state which contact counts are used to normalize the matrix, construct the two input distributions, estimate any distance–contact relationship and choose analysis settings. If the held-out contact counts influence any of these steps, the final comparison is not independent of the information used to construct the prediction.

**Measurement unit and test unit.** The unit being measured does not necessarily determine the unit that should be separated between training and testing. The split must follow the unit for which the outcome is evaluated. In lineage prediction, cells provide the early-state measurements, but the later outcome belongs to the lineage. All cells from one lineage must therefore remain together in either training or testing. Early measurements from a test lineage can be used to construct its predictors because those measurements are available at prediction time. Its later outcome remains withheld as the test target. A complete plan must specify both the lineage-level split and which time-specific measurements are available before prediction.

#### S2.3 What completion can add

Completion may add a missing specification only if the completed candidate remains the same analysis. The biological question, main result, analysis unit, transferred core method and established method–data correspondence must remain unchanged. Completion can write out a supported conversion, choose a value from an already allowed candidate set, specify how an intermediate quantity is calculated, or define which observations enter fitting and testing. Replacing response prediction with clustering changes the main result. Assigning a new biological meaning to an unsupported supply value changes the method–data correspondence. These changes define a different analysis and cannot be introduced through completion.

Completion can fill in a missing validation split, but it cannot repair a candidate that already defines a circular use of the test outcome. If the available observations support an independent split and the candidate simply omitted that split, completion can add it. If the candidate explicitly uses the final test outcome to choose inputs, relationships, representation or the main calculation, changing that procedure would define a different analysis.

When several incomplete candidates are eligible for completion, BioMINTREL first chooses the candidate with the fewest unmet completeness requirements. Ties are resolved using the combined and component scores defined in Section S2.4, followed by generation order. If no incomplete candidate can be completed without changing the analysis, the pair stops without a complete plan.

#### S2.4 Scoring and selecting completed plans

Completeness and plan scoring serve different purposes. Only candidates that pass every completeness requirement can enter final plan selection. The four scores do not determine whether a plan can be executed. They rank multiple complete plans for the same task–method pair according to four properties: how directly the required inputs are supported by the data, how clearly the transferred method produces the main result, how specifically the method–data correspondence is defined, and how fully the remaining analysis choices are fixed.

$g_D$ asks how directly the required inputs are supported by the data. $g_M$ asks whether the transferred method produces the main result and whether the route from method output to that result is fully specified. $g_C$ asks how completely and biologically specifically the method–data correspondence has been established. $g_A$ asks how fully the remaining material analysis choices have been fixed without using the final outcome. Table S1 lists the terms averaged in these four component scores and the relative ordering used for different construction types.

For scoring, directly observed inputs or existing annotations receive the highest support, followed by fixed constructions and then predefined finite choices. These preferences rank complete plans and resolve ties among candidates eligible for completion, and they do not determine whether a plan is complete.

A choice is material when different allowed values would change what is analysed, how the biological objects are related, what result is produced or which data are used to test that result. Examples include the graph definition, transport-cost formula or the rule that converts a method output into the primary biological quantity. Solver tolerance or iteration limits are not material choices when changing them leaves these scientific definitions unchanged.

**Table S1:** Terms averaged in the four scores used to rank complete plans. The final selection compares only candidates that pass every completeness requirement.

| **Score** | **Score components** | **Relative ordering** ($>$ means higher support) |
|---|---|---|
| $g_D$: input support | One construction term for each necessary input. | Direct observation / existing annotation $>$ fixed construction $>$ predefined finite choice. |
| $g_M$: core contribution | Three terms: how the core output becomes the primary result; whether the primary result actually requires the core calculation; and whether the route from core output to primary result is fully specified. | Native output $>$ fixed output conversion $>$ predefined finite output construction; satisfied $>$ uncertain $>$ violated. |
| $g_C$: correspondence | The construction term for every required input and relationship, plus one joint check that their biological meaning is supported and the method core remains preserved. | Input / relationship terms use the same construction order; joint check: satisfied $>$ uncertain $>$ violated. |

Supplementary Table S1 (continued)

| **Score** | **Score components** | **Relative ordering** (> means higher support) |
|---|---|---|
| $g_A$: analytical choices | Whether the calculation sequence is fixed, whether choices are independent of the final outcome, whether the stated definitions are sufficient to run the analysis, and how each material choice is specified. | Fixed calculation sequence / fixed material choice > predefined finite choice; general checks: satisfied > uncertain > violated. |

Different complete plans can contain different numbers of required inputs, relationships and material choices, so each component score averages over the terms relevant to that plan. For $g_C$, the denominator is the number of required inputs plus the number of required relationships plus one joint check of biological basis and core preservation. Thus, two inputs and one relationship give four terms. For $g_A$, the denominator is three general terms plus one term for each material choice. Two material choices therefore give five terms. Numerical solver settings that leave the scientific definitions unchanged do not add another material-choice term.

BioMINTREL uses two rankings for two different decisions. Before completion, it ranks incomplete eligible candidates to decide which candidate to complete. This ranking first minimizes the number of unmet completeness requirements, as described in Section S2.3. After completion, it ranks only complete candidates to choose the final plan. For this final ranking, the component scores and combined score are rounded to six decimal places. Equal combined scores are ordered by the larger minimum component score, then larger $g_C$, larger $g_A$, fewer completion attempts and earlier generation order.

**Supplementary Note S3: Details of the benchmark comparisons**

### S3.1 Defining benchmark tasks and reference labels

In BioMDC-Bench, a task is defined by both the data source and the starting biological question. The same source study therefore gives separate tasks when it supports different starting questions. D4/D5, D12/D13, D14/D15, D16/D17 and D18/D19 are such pairs of tasks. Each task is then paired with the same 12 candidate methods. Supplementary Data 1 (`Tasks; Methods; Pairs 228`) gives the task definitions, source articles, candidate methods and pair-level expert labels.

Each task–method pair has one fixed expert-consensus Tier that is used for every model comparison. The reference contains 24 T1, 152 T2 and 52 T3 pairs. The same reference label is used for direct judgment and BioMINTREL under both underlying language models.

### S3.2 Comparison between direct LLM judgment and BioMINTREL

The direct LLM baseline and the BioMINTREL correspondence condition receive the same task-specific and method-specific information for each pair. They differ only in how this information is converted into a Tier. The direct LLM baseline asks the language model to assign T1, T2 or T3 directly. In the BioMINTREL condition, the language model first constructs the method–data correspondence, and the fixed Tier rules then assign T1, T2 or T3 from that correspondence. Neither condition receives expert labels or retrieval scores. The comparison therefore tests whether explicitly constructing the method–data correspondence improves Tier judgment when the available pair information is held fixed.

The benchmark uses `gpt-5.4-mini-2026-03-17`[19] and `Qwen3.6-35B-A3B`[20]. For a given model, the direct LLM judgment and BioMINTREL correspondence calls use the same temperature of 0.1.

For each underlying language model, direct judgment and BioMINTREL produce judgments for the same 228 task–method pairs. The paired comparison therefore uses each pair as its own comparison unit. A "direct-only correct" pair agrees with the expert reference only under direct judgment, whereas a "BioMINTREL-only correct" pair agrees only after the correspondence workflow. The paired test compares these two discordant counts and asks whether one condition agrees with the expert reference more often on the same pairs.

We also ask whether BioMINTREL gives the same Tier judgment when the underlying language model is changed. We run BioMINTREL separately with GPT-5.4 mini and Qwen3.6-35B-A3B and compare their T1+T2 versus T3 assignments for the same task–method pairs. It measures how consistently BioMINTREL classifies the same pairs across the two underlying language models. The pairwise joint counts can be derived from Supplementary Data 1 (`Model decision`).

### S3.3 Later-literature labels

A positive later-literature label means that the literature collection contains at least one study published in 2021 or later that uses the same or a closely related analytical approach for the target task or a neighbouring biological question. The later study does not need to reproduce BioMINTREL's complete correspondence or final analysis plan. The label therefore records later research support for the task–method pairing.

Examples of matches under this rule include PASTE and PASTE2 for spatial alignment with Gromov–Wasserstein methods

(D4/M2)[65, 66], collective optimal transport for spatial communication (D19/M1)[27], and PertSpectra for matrix factorization in perturbation-response tasks (D8/M3 and D10/M3)[67]. Supplementary Data 1 (`Pairs 228`) gives all pair-level mappings and study identifiers.

Each pair contributes one binary literature outcome. One or more matching studies gives a positive record, and additional studies do not give that pair additional weight. A single study can support more than one task–method pair when it provides evidence relevant to several combinations. "No related study found" means that no study in the collection met the matching rule for that pair.

Task-level averages and pooled proportions answer different questions. The task-level average evaluates how consistently later-literature support appears across biological tasks. For each task, we calculate the proportion of literature-supported pairs separately within its predicted T1+T2 and T3 groups, and then give each task with a defined proportion equal weight. If a task contains no pair in a given predicted group, that proportion is undefined and is omitted from the average. A task with one pair in the group and a task with more than one pair in the group therefore contribute equally at the task level.

The pooled proportion evaluates the overall later-literature support among all pairs assigned to that group. It divides the total number of literature-supported pairs by the total number of pairs in the group across all tasks, so tasks contributing more pairs to that group receive more weight. Reporting both quantities shows whether the pattern is consistent across biological tasks and how strong the overall pair-level literature support is. The underlying pair-level records are in Supplementary Data 1 (`Pairs 228`).

#### S3.4 Human and model judgments on later-supported pairs

Among the 228 BioMDC-Bench pairs, 170 have a positive later-literature label, meaning that a related application was found in studies published in 2021 or later. On these 170 pairs, we compare the Tier assignments from the expert reference and the two BioMINTREL implementations. We combine T1 and T2 because both indicate that the method can be applied, either directly or with defined adaptations. Coverage is the proportion of the 170 pairs assigned to T1+T2. This analysis asks how often each judgment source recognizes an application among pairs later supported by literature. The analysis of all 228 pairs asks a different question: whether T1+T2 pairs are more likely than T3 pairs to receive later-literature support (Section S3.3). See Supplementary Data 1 (`Pairs 228; Model decision`) for the original Tiers and literature records.

Coverage and binary agreement answer two different questions. Coverage asks how many of the 170 later-supported pairs each source assigns to T1+T2. Binary agreement asks whether experts and BioMINTREL make the same T1+T2-versus-T3 decision for each individual pair. For GPT-5.4 mini, the joint counts are 133 pairs assigned to T1+T2 by both sources, 16 by experts alone, 11 by BioMINTREL alone and 10 assigned to T3 by both. For Qwen3.6-35B-A3B, the corresponding counts are 117, 32, 17 and 4. Binary agreement therefore equals 143/170 (84.1%) and 121/170 (71.2%). The three-level cross-tabulations retain the distinction between direct application and defined adaptation (Supplementary Data 1 (`Pairs 228; Model decision`)).

Because the same 170 pairs are judged by experts and BioMINTREL, we compare their T1+T2 coverage with two-sided exact McNemar tests[57, 58] and apply Holm correction[61] to the two human–model comparisons. The adjusted values are 0.442068 and 0.088768.

#### S3.5 Comparison between scientific context and analytical structure retrieval

The retrieval stress test is defined from expert T2 pairs using Scientific context similarity only. For each BioMINTREL implementation in `gpt-5.4-mini-2026-03-17` and `Qwen3.6-35B-A3B`, the 38 T2 pairs with the lowest Scientific context similarity form that implementation's evaluation subset. The two implementations select their subsets separately because their Scientific context scores are obtained from their own runs.

After the subset is fixed, Scientific context and Analytical structure are compared on exactly the same selected pairs within that implementation. For each selected pair $(t_i, m_j)$, the target method $m_j$ is ranked against the 12 candidate methods for the same task $t_i$. Tied methods receive their average occupied rank. For example, a tie spanning positions 3 and 4 gives rank 3.5. A change in rank therefore reflects the retrieval representation used for the same task and candidate-method set.

#### S3.6 Prioritization by scientific criteria

This step tests whether the three scientific criteria improve the choice of methods that advance to plan generation. Each task can send only three methods forward. We therefore compare the expert-reference support among the top three methods when candidates are ranked by Tier alone and when the scientific criteria are added.

For the evaluation, we convert Tier and scientific-criteria issues into an ordinal score. T1 starts at 30, T2 at 20 and T3 at 0. Each identified scientific-criteria issue subtracts one point, with a maximum deduction of two points in this evaluation. The resulting scores are:

$$\begin{aligned}
\text{T1}: &\quad 30,\ 29,\ 28 \quad \text{for 0, 1, or 2+ issues,}\\
\text{T2}: &\quad 20,\ 19,\ 18 \quad \text{for 0, 1, or 2+ issues,}\\
\text{T3}: &\quad 0.
\end{aligned}$$

These values encode candidate order only. The ten-point gap keeps every T1 candidate ahead of every T2 candidate, while the issue count changes the order within the same Tier. In the Tier-only condition, T1, T2 and T3 receive scores of 2, 1 and 0. Candidates within the same Tier therefore tie, so this condition measures the prioritization provided by Tier alone.

Fig. 2e evaluates the top three positions for each task. When a tie crosses the third-place boundary, the evaluation assigns the remaining place according to the fraction of tied methods that are T1+T2 in the expert reference. For example, if one place remains and two of four tied methods are T1+T2 in the expert reference, that place contributes 0.5 expected supported methods. This prevents an arbitrary method identifier from changing the evaluation result.

The actual pipeline must select three specific methods for plan generation. It first orders candidates by Tier and then uses the full number of scientific-criteria issues within the same Tier. If candidates still tie, the fixed pair identifier determines their order. The full issue count remains available here, so candidates with two and three issues can still receive different priorities.

The Fig. 2e evaluation score measures how much the scientific criteria improve top-three expert-reference support. The pipeline selection rule determines which three methods actually proceed to plan generation.

#### S3.7 Independent plan generations

This evaluation tests whether repeated independent generation increases the chance that the same task–method pair produces at least one complete analysis plan. For each selected pair, BioMINTREL generates five initial plans independently from the same task, method and correspondence information. Each generation is produced without access to the other candidate plans. Fig. 2f uses only these five independent initial plans, so it isolates the contribution of repeated formulation itself.

For a condition allowing $n$ generations, a pair is counted as complete when at least one of its first $n$ initial plans satisfies all completeness requirements. For example, if the first two plans are incomplete and the third plan is the first complete one, the pair is incomplete in the one- and two-generation conditions and complete in the three-, four- and five-generation conditions. A blank entry means that none of the five independent initial plans was complete. For each underlying language model, the same 57 selected pairs are used in all one- through five-generation conditions. Fig. 2f therefore shows how many selected pairs have produced at least one complete plan as the number of allowed independent generations increases.

Confidence intervals are obtained by bootstrapping biological tasks[62]. Each task contributes three selected method pairs that share the same data and starting biological question. When a task is sampled into a bootstrap replicate, all three of its selected pairs enter together. If the same task is sampled twice, all three pairs are repeated twice. This preserves the grouping of method pairs within a biological task and makes the interval reflect variation across tasks.

### Supplementary Note S4: Expert review criteria and summary rules

#### S4.1 Plan labels and scoring boundaries

The anonymous letters do not represent the same plan type across all four cases. Reviewers assessed the candidates under the anonymous labels A, B and C. The full plan descriptions are provided in Supplementary Data 3 (`plan-review forms`).

The five 1–4 scores evaluate the plan itself: novelty relative to the source paper, biological relevance, methodological reliability, transfer fit and validation sufficiency. Reviewer confidence is scored separately on a 1–3 scale and records how certain the reviewer is about their own assessment. The proceed question is a separate yes/no judgment of whether the reviewer would pursue that plan in their own group. The original plan-review forms provide the questions and score anchors (Supplementary Data 3 (`plan-review forms`)).

#### S4.2 What the three plan-level choices measure

The overall ranking, the additional-information choice and the proceed recommendation answer three different questions and are counted separately. The overall ranking compares the three plans within the same case and asks which plan is most worth executing. The additional-information question asks which single plan is most likely to provide genuinely additional information beyond the analyses already available for that case. The proceed question is answered separately for each plan and asks whether that plan is worth pursuing. A reviewer can therefore recommend more than one plan to proceed while selecting only one plan as most likely to provide additional information.

Each plan type receives eight score assessments: two reviewers across four cases. The additional-information question also gives eight choices in total, one for each reviewer–case assessment. Proceed is a separate yes/no decision for each individual plan, so each plan type receives eight proceed decisions. First-place counts can exceed eight in total because ties are allowed. In Case 4, Reviewer 1 ranked plans B and C jointly first, so both plans receive one first-place count. The eight reviewer–case assessments therefore produce nine first-place assignments. Individual scores, rankings, choices and comments are provided in Supplementary Data 3 (`plan-review forms`).

#### S4.3 Scoring the completed analyses

The completed-analysis review evaluates each BioMINTREL case after its results and validation are available. The five 1–4 scores assess biological relevance, framework–question fit, methodological soundness, evidence and validation, and information gain relative to a conventional target-only analysis. Reviewer confidence again records how certain the reviewer is about their own assessment and uses a separate 1–3 scale. The follow-up question is a separate yes/no judgment of whether the completed analysis is worth further study. The original completed-analysis review forms provide the questions and score anchors (Supplementary Data 3 (`completed-analysis review forms`)).

Individual completed-analysis scores and open comments are provided in Supplementary Data 3 (`completed-analysis review forms`).

### Supplementary Note S5: PjDHFR responses across methotrexate doses

**S5.1 Analysis populations.** The analysis compares the effects of amino acid substitutions in *Pneumocystis jirovecii* dihydrofolate reductase (PjDHFR) across methotrexate doses. The source study measures coefficients of selection (CoS) in a dimethyl sulfoxide control (DMSO) and lower- and higher-dose methotrexate conditions, MTX4 and MTX20 [21]. Each condition contrast uses all records with both CoS measurements and their standard errors. Response classes and position enrichment use single-amino-acid substitutions. The latent-phenotype fit includes synonymous and wild-type records and excludes stop variants. These populations match the information required by each analysis.

For sequence fitting, each substitution changes its stated position in the wild-type sequence. Synonymous and wild-type records share the unchanged sequence. Structural comparisons additionally require an available FoldX score. This requirement retains 101 of the 102 high-dose-specific substitutions and leaves their response-class assignments unchanged.

**S5.2 Response classes across doses.** Condition contrasts separate drug responses from the DMSO control and compare the two drug doses. Each contrast subtracts the two CoS values and combines their standard errors as the square root of the sum of their squares. A zero combined standard error leaves the test value undefined. Two-sided normal tests and Benjamini–Hochberg adjustment apply separately to the MTX4–DMSO, MTX20–DMSO and MTX20–MTX4 comparisons [68]. Each contrast therefore defines its own set of tests for adjustment.

The response classes combine the direction of selection under each drug condition with evidence for a difference from DMSO. In Table S2, $c_D, c_4, c_{20}$ denote the CoS values themselves, and $q_4, q_{20}$ denote the adjusted probabilities for the two drug–DMSO contrasts. The sign conditions apply to $c_4$ and $c_{20}$. The separate MTX20–MTX4 contrast provides a dose comparison and does not determine class membership. Each substitution receives the first class whose conditions it satisfies.

**Table S2:** Sequential definitions of the four response classes. Each record receives the first class whose conditions it satisfies.

| Class | Conditions |
|---|---|
| Cross-dose resistant | $c_4 > 0,\ q_4 < 0.01,\ c_{20} > 0,\ q_{20} < 0.01,\ \lvert c_D \rvert < 0.1.$ |
| High-dose-specific | $c_{20} > 0,\ q_{20} < 0.01,\ \lvert c_D \rvert < 0.1$, and at least one of $q_4$ undefined, $q_4 \geq 0.01$, or $c_4 \leq 0$. |
| Drug-sensitive | $c_4 < 0$ with $q_4 < 0.01$, or $c_{20} < 0$ with $q_{20} < 0.01$. |
| Neutral | All remaining records. |

The sequential rule gives each substitution one response class. For example, a substitution with negative MTX4 CoS, positive MTX20 CoS and significant contrasts against DMSO at both doses enters the high-dose-specific class when $\lvert c_D \rvert < 0.1$, even though it also meets the drug-sensitive criterion. The neutral class contains the substitutions remaining after the first three rules.

**S5.3 Positional and structural enrichment.** Positional enrichment asks whether a response class concentrates at particular residues. Its frequency among all 3,762 substitutions supplies the background. At each position, a two-sided binomial test compares the observed class count with this background, using the number of tested substitutions at that position as the denominator. This comparison accommodates differences in coverage across positions. Benjamini–Hochberg adjustment applies separately across positions within each class.

The source FoldX score $\Delta\Delta G_{\mathrm{MTX}}$ describes the substitution-associated energy difference in the methotrexate-bound protein model[21, 69]. Structural comparisons use records with a defined score. Kruskal–Wallis tests compare the four response classes and, separately, positions G124, G125 and G126[70]. Two-sided Mann–Whitney tests compare all six class pairs[71], with Benjamini–Hochberg adjustment across those six tests.

**S5.4 Latent phenotype and residual candidates.** MAVE-NN learns an additive map from amino acid sequence to a one-dimensional latent phenotype $\phi$, with a linear measurement function and Gaussian observation noise [23]. Fitting uses 60 epochs, batches of 512, the Adam optimizer[72] at learning rate $10^{-3}$, shuffled input order and a 20% internal validation fraction. Calibration then places $\phi$ on the MTX20 CoS scale, so a residual measures the difference between an observed response and the calibrated sequence-based expectation.

Calibration divides the substitution records into 30 equal-frequency bins of $\phi$, removing duplicate boundaries. Each bin contributes its midpoint $(\min\phi+\max\phi)/2$ and its median MTX20 CoS. Linear interpolation between these points defines the expected response $\hat{c}_{20}(\phi)$. Only records within the bin-midpoint range receive a calibrated residual, and calibration uses no extrapolation.

The residual is $r = c_{20} - \hat{c}_{20}(\phi)$. Its standard deviation, $\mathrm{SD}(r)$, uses all non-stop records with a defined residual, including synonymous and wild-type records. The strict thresholds $r > 2\,\mathrm{SD}(r)$ and $r < -2\,\mathrm{SD}(r)$ define the high and low residual sets. Response-class composition then uses the single-amino-acid substitutions within each set. This retains a common residual threshold while matching the biological summary to the substitution population.

Residual enrichment asks whether these departures from the sequence-based expectation favour particular response classes. A one-sided Fisher test[60] compares membership in a response class between one residual set and all other substitutions in the 3,762-substitution population. Class composition, enrichment and recurrent positions identify residual candidates for further testing.

**Supplementary Note S6: Spatial constraints on ligand receptor communication**

**S6.1 Spatial graph and expression scale.** The spatial analysis asks how ligand-associated supply and receptor-associated capacity constrain possible communication in a breast-cancer tissue section [24]. The Visium data provide expression counts and full-resolution coordinates for 1,657 tissue locations, each represented by one graph node. Expression normalization to a total count of $10^4$ per location, followed by $\log(1 + x)$, defines the scale used for allocation.

A directed graph connects each location to its ten nearest other locations by Euclidean distance. These 16,570 connections exclude self-connections. Retaining distances at or below their 95th percentile, 730.03 coordinate units, leaves 15,806 connections. For each ligand–receptor pair, candidate connections link ligand-positive senders to receptor-positive receivers. Each pair therefore has its own expression-eligible graph within the same spatial neighbourhood structure.

**S6.2 Supply, capacity and allocation costs.** Supply represents ligand-associated expression available for allocation, and capacity limits how much receptor-associated expression can receive. For a sender $i$ and receiver $j$, $L_i$ and $R_j$ are ligand and receptor expression divided by their respective tissue-wide maxima. Integer supply and capacity are $s_i = \mathrm{round}(100L_i)$ and $c_j = \mathrm{round}(100R_j)$, using nearest-even rounding. These model units encode normalized expression, and they do not measure molecule counts. The tissue maximum maps to 100 before rounding, while a location with positive expression can have zero integer supply or capacity.

An allocation cost combines the expression available at both ends of a connection with their spatial separation. For a candidate connection $(i, j)$, let $d_{ij}$ be its distance and $d_{\max}$ the largest distance among that pair's expression-eligible connections. The unit cost is

$$a_{ij} = \mathrm{round}\left[1000\left\{1 - \left(0.4L_i + 0.4R_j + 0.2\left(1 - \frac{d_{ij}}{d_{\max}}\right)\right)\right\}\right].$$

Higher sender expression, higher receiver expression and shorter distance each lower the cost. Both normalized distance and the combined score are clipped to $[0, 1]$; normalized distance is zero if $d_{\max} = 0$. Each connection has capacity $\min(s_i, c_j)$.

**Allocation objective.** Let $x_{ij}$ be allocated integer flow, $u_i$ unallocated sender supply and $\mathcal{E}$ the candidate connection set. Each pair solves [25, 26]

$$\begin{aligned}
\min_{x,u} \quad & \sum_{(i,j)\in\mathcal{E}} a_{ij}x_{ij} + 1200\sum_i u_i,\\
\text{subject to} \quad & \sum_{j:(i,j)\in\mathcal{E}} x_{ij} + u_i = s_i,\\
& \sum_{i:(i,j)\in\mathcal{E}} x_{ij} \le c_j,\\
& 0 \le x_{ij} \le \min(s_i, c_j), \qquad u_i \ge 0.
\end{aligned}$$

The supply constraint assigns every unit to a permitted connection or leaves it unallocated. Receivers share their capacity across incoming connections. Unallocated supply costs 1200 per unit and preserves the amount that spatial connections or receiving capacity cannot accommodate.

**S6.3 Allocation summaries.** Supply fractions describe how much the network allocates, while the sender fraction describes how widely unallocated supply occurs across the tissue. The allocated and unallocated fractions both divide by total sender supply. The fraction of senders retaining supply divides the number of ligand-positive locations with any unallocated supply by all ligand-positive locations.

Receiver utilization is $\sum_i x_{ij}/c_j$ for $c_j > 0$. Mean utilization averages receivers with positive integer capacity. A receiver reaches capacity when its utilization is at least 0.99. The proportion at capacity divides that count by all receptor-positive locations, including those whose scaled capacity rounds to zero. Mean utilization and the proportion at capacity thus describe different receiver sets.

Candidate degree describes the connections permitted by spatial and expression constraints. Used degree describes connections carrying positive flow. Mean incoming degree averages all 1,657 locations, including zeros. The used-connection fraction divides positive-flow connections by candidate connections for the same ligand–receptor pair. Together, these quantities describe how allocation concentrates flow within the available network.

**S6.4 Method comparisons and spatial context.** Comparisons with established communication methods match quantities at the spatial level each method provides. COMMOT uses the same transformed expression matrix and 1,657 locations, an exact-pair ligand–receptor database and a distance threshold of 730.0 coordinate units [27]. Its sender and receiver scores match outgoing and incoming allocation for the same pair and location. CellChat supplies communication between Leiden groups at the pathway level, with a 250-micrometre spatial interaction range [28, 73]. Its pathway scores provide a separate comparison at that level.

Pearson and Spearman correlations compare COMMOT scores with allocated flow across all locations. Marker enrichment examines the tissue context of receiving hotspots. These hotspots comprise receptor-positive locations with incoming flow at or above the 90th percentile, including ties. One-sided Mann–Whitney tests compare each of eight marker genes between hotspots and all other tissue locations. Benjamini–Hochberg adjustment covers the eight genes within each ligand–receptor pair[68].

The TGFB1–TGFBR1 distance summary measures each ligand-positive location's distance to its nearest receptor-positive location. Coexpression at the same location gives distance zero. This summary describes local receptor availability. Allocation follows the graph's exclusion of self-connections, so coexpression contributes to the distance summary without creating a self-allocation edge.

**Local display and tissue-wide comparisons.** Fig. 4b shows a common square region with half-width 3,500 full-resolution coordinate units for all five ligand–receptor pairs. The selected region maximizes the sum, across the five pairs, of the fraction of each pair's whole-tissue flow contained in the region. A common region allows the displayed patterns to share the same tissue context. The figure draws the 45 strongest connections within it, and node flow includes all connections in that region. All quantitative comparisons use the complete tissue.

**Supplementary Note S7: Perturbation response paths through cell states**

**S7.1 Shared transcriptional state space.** The unfolded protein response (UPR) experiment provides a common transcriptional state space for comparing responses to genetic perturbations [37]. Cells express at least 200 genes and have at most 40% mitochondrial counts, while genes occur in at least three cells. The expression representation combines normalization to 10,000 counts per cell and $\log(1 + x)$ transformation with 3,000 highly variable genes selected from counts, gene scaling clipped at 10, 30 principal components and 20 nearest neighbours. Leiden clustering[74] at resolution 0.6 and seed 0 defines the same 11 states for all perturbations.

A programme score is the mean normalized, log-transformed expression of the available genes in a predefined set. The source defines 37 genes for protein kinase R-like endoplasmic reticulum kinase (PERK) and the integrated stress response, 19 for activating transcription factor 6 (ATF6) and chaperones, and 21 for inositol-requiring enzyme 1 (IRE1), X-box binding protein 1 (XBP1) and the translocon. Standardizing each branch score across cells by its population standard deviation and averaging the three standardized scores gives total UPR. The shared induced-response and high-abundance-response sets contain 20 and 12 available genes. Each state's programme score averages its cells' scores.

Each perturbation defines one query. The target list contains 17 perturbations, and the expression data assign cells to 15 of them. GARS and WARS have no assigned cells. Summing cell-neighbour connection weights between states gives a shared undirected graph with 55 edges. Keeping this state space and its underlying connections fixed allows each query's boundary states and directed weights to express its particular response.

**S7.2 Response direction and boundary states.** Boundary states define where a modelled response route begins and ends. For state $a$, the control fraction $C_a$ and selected-perturbation fraction $Q_a$ both use the state's total cell count $N_a$ as denominator. Let $B_a$ denote the relevant branch score and $U_a$ total UPR. The source annotation assigns the IRE1/XBP1/translocon score to $B_a$ for IRE1-selective perturbations and total UPR for the other selected response patterns.

The notation $z(\cdot)$ denotes standardization across the 11 states by the population standard deviation. The starting score $s_a$, target score $t_a$ and progress score $h_a$ are

$$
\begin{aligned}
s_a &= 1.25z(C_a) - 0.90z(Q_a) - 0.50z(B_a) - 0.25z(U_a) + 0.20z(\log(1 + N_a)),\\
t_a &= 1.25z(Q_a) + 0.75z(B_a) + 0.45z(U_a) - 0.30z(C_a) + 0.15z(\log(1 + N_a)),\\
h_a &= 1.10z(Q_a) + 0.70z(B_a) + 0.35z(U_a) - 0.70z(C_a).
\end{aligned}
$$

The starting and target scores select boundary states, while progress defines the preference for movement between them. Control representation raises the starting score. Perturbation representation and response raise the target and progress scores.

Among states with at least 40 cells, the two highest starting scores and two highest target scores define the initial boundary sets. Ties favour control fraction for starts and perturbation fraction for targets, followed by state size. A state shared by both sets retains starting membership and leaves the target set. If no target remains, the highest-ranked non-starting state becomes the target. This rule keeps the boundaries disjoint and allows the final target set to contain fewer than two states.

Progress orients the common graph through $W_{ab} = w_{ab} \exp[(h_b - h_a)/2]$, where $w_{ab}$ is the shared edge weight. When $h_b > h_a$, the weighting favours movement from $a$ to $b$. The reverse direction uses the opposite difference. A self-connection contributes 0.05 times each state's total outgoing weight to other states. Row normalization then gives the perturbation-specific transition probabilities $P$.

**S7.3 Route probabilities and gateway states.** The forward committor $q_a^+$ is the probability of reaching the target set before the starting set under transition matrix $P$. It is 0 in the starting set and 1 in the target set. Intermediate states satisfy $q_a^+ = \sum_b P_{ab} q_b^+$ [38, 75]. The backward probability $q_a^-$ describes whether a trajectory's most recent boundary visit was to the starting set. The stationary weight $\pi_a$ is the long-run probability of state $a$ under this transition matrix. $\pi P = \pi$ and $\sum_a \pi_a = 1$. The time-reversed matrix is $P_{ab}^{\text{rev}} = \pi_b P_{ba}/\pi_a$. Its boundary problem sets $q^-$ to 1 in the starting set and 0 in the target set[76].

Gross reactive flux, $f_{ab} = \pi_a q_a^- P_{ab} q_b^+$, weights the contribution of a directed connection to routes linking the boundaries. Net flux retains the positive difference between opposing contributions, $F_{ab} = \max(f_{ab} - f_{ba}, 0)$[75, 76]. For an intermediate state,

$$G_a = \min\left(\sum_b F_{ba}, \sum_b F_{ab}\right).$$

The smaller of incoming and outgoing net flux determines $G_a$, so a high score requires flow to enter and continue through a state. The three intermediate states with the highest $G_a$ values form the gateway set. Starting and target states remain outside this ranking.

Gateway membership describes passage through an intermediate state, while arrival probability describes access to the target set. Targets have arrival probability 1 and do not compete for gateway membership. Gateway recurrence counts the perturbations that include a state among their three selected intermediates. Target recurrence counts those that select it as a target. Both counts use the same 15 perturbations.

Pearson correlations compare forward-probability vectors across perturbations using the same 11 states. Cosine similarity compares net-flux vectors on one common list of directed connections, assigning zero to a connection with no flux for a query. Shared state and connection coordinates make these comparisons reflect changes in route probabilities and flow across the same transcriptional landscape.

**S7.4 Null model for recurrent gateways.** The null comparisons ask whether a state recurs as a target or gateway more often than expected from the query composition and the common state graph. The selected-query comparison redistributes the 15 perturbation labels among their pooled cells while retaining each perturbation's cell count. The all-non-control comparison draws groups of those same sizes from all non-control cells. Controls retain their labels. Each comparison uses 5,000 permutations, testing recurrence against the selected responses and against the broader perturbation pool.

Boundary selection belongs to the procedure under test because changes in each state's perturbation fraction can change its starting and target roles. Each permutation therefore repeats boundary selection and overlap removal, progress-based graph orientation, and the calculation of arrival probabilities, net flux and gateways. Expression states, undirected connections and programme means remain fixed. This null tests recurrence after the same boundary selection used in the observed analysis.

A state-specific comparison places the observed recurrence against that state's recurrence in every permutation. Adjustment for searching across states uses the largest recurrence among all 11 states in each permutation, with separate maxima for target and gateway recurrence[77]. These comparisons address evidence for a specified state and evidence after selecting a state from the full set, respectively.

**Supplementary Note S8: Rare cell states and sampling within lineages**

**S8.1 Programme scores and lineage populations.** The Watermelon data connect early transcriptional states with later abundance in PC9 lung-cancer lineages [45]. Prediction uses the 2,192 lineage identifiers present on both days 3 and 14: day-3 cells supply the features and the day-14 count supplies the outcome. Matching the two time points defines the population for this question.

A gene-programme score summarizes a cell's expression of a specified gene set. The candidate programmes are xenobiotic metabolism, glutathione metabolism and reactive oxygen species (ROS) defence. Notch and interferon-alpha provide biological controls. Their selection follows evidence on rare transcriptional states and drug resistance [49, 50] and pathway findings in the source study [45]. The analysis uses the existing cell-level programme and cell-cycle scores.

**Residualization.** Within each time point, ordinary least squares adjusts each programme score for the available cell-cycle score, unique molecular identifier count, detected gene count and mitochondrial percentage. Each fit includes an intercept, uses complete observations, omits constant covariates and requires at least ten cells. Residuals supply the adjusted cell scores. Fitting these adjustments to day-3 cells defines the early predictors independently of the day-14 lineage outcome.

**S8.2 Reading the upper tail.** The upper tail describes cells with high programme scores within a lineage. Let $n$ count the available scores and $q$ denote the quantile level. At $q = 0.95$, the top-5% mean uses one cell when $n = 10$ and two cells when $n = 21$. The same nominal quantile can therefore represent a maximum in one lineage and an average of several high-scoring cells in another. The analysis orders signed residual scores from low to high, so the upper tail captures high programme activity after adjustment.

**Empirical tail estimator.** The analysis draws on conditional value at risk, a summary of a specified distributional tail [46–48]. For one lineage and programme, order the $n$ available adjusted scores as $s_{(1)} \leq \cdots \leq s_{(n)}$. At quantile $q$, the number of included cells and their mean are

$$k_q = \max\{1, \lceil n(1-q) \rceil\}, \qquad T_q = \frac{1}{k_q} \sum_{r=n-k_q+1}^{n} s_{(r)}.$$

Here $\lceil \cdot \rceil$ rounds upward. The primary analysis uses $q = 0.95$. When $k_q = 1$, $T_q$ equals the largest observed score. Larger $k_q$ averages several high-scoring cells. Missing scores enter neither the tail nor the lineage mean, which averages all available scores. A lineage without available scores has no summary.

**Sampling depth.** Fig. 6a,b describes all 36,239 day-3 lineage–score records from 5,177 lineages across seven scores: the five prediction programmes, fatty acid metabolism and the cell-cycle score. This population establishes the sampling depth available for an upper-tail summary. Prediction additionally requires a day-14 outcome and uses the 2,192 matched lineages in Section S8.1. The sampling and prediction populations therefore have distinct denominators.

**S8.3 Prediction models.** For lineage $\ell$, let $N_{\ell,t}$ denote the sampled cell count on day $t$. The early-abundance predictor is $a_\ell = \log(1 + N_{\ell,3})$, and the outcome is $y_\ell = \log(1 + N_{\ell,14})$. The day-14 count pools cells carrying the same lineage barcode across both experimental replicates and the three mCherry sorting fractions, measuring lineage abundance among sampled cells. Let $m_\ell$ and $T_\ell$ denote the day-3 mean and top-5% score. Linear models with an intercept use $m$, $T$, $a$, $(a, m)$ or $(a, m, T)$. Comparing $(a, m, T)$ with $(a, m)$ asks whether the upper tail adds predictive information while retaining the same abundance and mean predictors.

Ten-fold cross-validation repeats ten times with base seed 20260618. All models for a programme share the same lineage divisions, with each lineage entirely in one fold. Test-fold $R^2$ uses that fold's outcome mean as reference. Averaging the 100 scores gives model performance. Averaging the matched fold-wise differences between $(a, m, T)$ and $(a, m)$ gives the upper-tail increment. Matching folds keeps the model comparison on the same held-out lineages.

**S8.4 Exact-size permutations.** Exact-size permutations test the association between early state summaries and later abundance at a fixed sampling depth. The paired summaries $(m_\ell, T_\ell)$ move together between lineages with identical day-3 cell counts, while each lineage retains its early abundance and later outcome. Joint movement preserves the mean–tail correlation, and restriction to equal cell counts preserves the sampling-depth distribution. Each of 500 permutations refits $(a, m)$ and $(a, m, T)$ and recalculates their cross-validated difference.

The observed permutation statistic and all 500 permuted statistics use one fixed ten-fold lineage division for each programme. This matching defines the null comparison on common folds. The ten repeated divisions in Section S8.3 summarize predictive performance. The test uses the upper tail of the permutation distribution[77], with Benjamini–Hochberg adjustment across the five programmes[68].

**S8.5 Equal-cell and minimum-size comparisons.** Equal-cell sampling compares lineages at the same observed depth. At each depth of 2, 3, 5 or 10 day-3 cells, eligible lineages contribute that number of available cells without replacement. Each draw supplies the same sampled cells to all five programmes and defines new mean and tail summaries. The analysis repeats the draw 100 times. Higher depths also restrict the eligible lineage population.

Within each draw, a fixed five-fold lineage division compares a mean-only model with a mean-plus-tail model. Equal sampled cell counts allow this comparison to omit abundance and isolate the contribution of the tail beyond the mean. The full-data comparison in Section S8.3 retains abundance in both models because lineage sizes vary there.

At depths of 2, 3, 5 and 10 cells, the top-5% summary contains one cell. The equal-depth comparison therefore tests whether the sample maximum adds predictive information beyond the lineage mean when every lineage contributes the same number of cells. The mean increment and its 2.5th–97.5th percentile range across the 100 draws summarize sensitivity to which cells enter each sample.

**Quantile and minimum-count sensitivity.** Quantile sensitivity compares $q$ values of 0.90, 0.95 and 0.99 in the same 2,192 matched lineages. Minimum-count sensitivity requires at least 1, 2, 3, 5, 10 or 20 day-3 cells and recalculates features in each eligible population. Both compare $(a, m, T)$ with $(a, m)$ over ten repeated divisions. A setting requires at least 20 lineages. Populations with at least 100 lineages use ten folds, and smaller eligible populations use five. The minimum-count analysis uses a setting-specific seed, which accounts for numerical differences between its one-cell row and the $q = 0.95$ quantile analysis despite their identical lineage populations. These analyses distinguish the choice of tail fraction from the choice of eligible population.
